# A consensus-based and readable extension of *Li*near *Co*de for *R*eaction *R*ules (LiCoRR)

**DOI:** 10.1101/2020.05.31.126623

**Authors:** Benjamin P. Kellman, Yujie Zhang, Emma Logomasini, Eric Meinhardt, Austin W. T. Chiang, James T. Sorrentino, Chenguang Liang, Bokan Bao, Yusen Zhou, Sachiko Akase, Isami Sogabe, Thukaa Kouka, Iain B.H. Wilson, Matthew P. Campbell, Sriram Neelamegham, Frederick J. Krambeck, Kiyoko F. Aoki-Kinoshita, Nathan E. Lewis

**Affiliations:** Department of Pediatrics, University of California San Diego School of Medicine, La Jolla, California, USA; Department of Bioengineering, University of California San Diego School of Engineering, La Jolla, California, USA; Bioinformatics and Systems Biology Program, University of California San Diego School of Engineering, La Jolla, California, USA; Department of Linguistics, University of California San Diego, La Jolla, California, USA; Graduate School of Engineering, Soka University, Hachioji, Tokyo, Japan; Faculty of Science and Engineering, Soka University, Hachioji, Tokyo, Japan; Department of Chemical and Biomolecular Engineering, Johns Hopkins University, Baltimore, Maryland, United States of America; ReacTech Inc., Alexandria, Virginia, United States of America; Institute for Glycomics, Griffith University, Gold Coast, QLD, Australia; Systems Glycobiology Consortium; Novo Nordisk Foundation Center for Biosustainability at the University of California San Diego School of Medicine, La Jolla, California, USA; Institute of Biochemistry, University of Natural Resources and Life Sciences, Vienna; Department of Chemical and Biological Engineering School of Engineering and Applied Sciences, State University of New York, University at Buffalo, Buffalo, New York, USA

**Keywords:** Systems glycobiology, Linear Code, Glycoinformatics

## Abstract

Systems glycobiology aims to provide models and analysis tools that account for the biosynthesis, regulation, and interactions with glycoconjugates. To facilitate these methods, there is a need for a clear glycan representation accessible to both computers and humans. Linear Code, a linearized and readily parsable glycan structure representation, is such a language. For this reason, Linear Code was adapted to represent reaction rules, but the syntax has drifted from its original description to accommodate new and originally unforeseen challenges. Here, we delineate the consensuses and inconsistencies that have arisen through this adaptation. We recommend options for a consensus-based extension of Linear Code that can be used for reaction rule specification going forward. Through this extension and specification of Linear Code to reaction rules, we aim to minimize inconsistent symbology thereby making glycan database queries easier. With a clear guide for generating reaction rule descriptions, glycan synthesis models will be more interoperable and reproducible thereby moving glycoinformatics closer to compliance with FAIR standards. Reaction rule-extended Linear Code is an unambiguous representation for describing glycosylation reactions in both literature and code.

## Introduction

Glycans are predominantly synthesized through the serial addition of monosaccharides to form large polysaccharides. To build computational models of glycan synthesis, the biochemical reactions involved must be defined and described mathematically in a form that can be interpreted by computers^1–3^. Several groups have created such models using a variety of strategies, including mechanistic and nonlinear^4–12^, linear probabilistic^13,14^, machine learning^15^, formal-grammar^16^ and substructural^17^. Unfortunately, most of these approaches use slightly different expressions of the building blocks, the reaction rules, therefore, model comparison is extremely challenging.

In the past few decades, substantial efforts made in the construction of these models of glycan synthesis were mostly focused on defining reaction rules that benefit from an unambiguous representation with human readability. For example, graphical denotation is one of the most human-understandable representations to describe reaction rules^18–20^. While graphical representations are intuitive and extremely accessible to a human reader, they are not computationally accessible due to ambiguities in their representations. There are already efforts to create computationally transmissible rule sets in XML and SBML ^7,8^ which are readily interoperable and reusable. However, the XML and SBML representations can be less human-readable, making them hard to include in the main text of a manuscript. As systems glycobiology develops, there is a need to develop a standard nomenclature for unambiguous and readable reaction rules to facilitate development, exchange, extension, and validation of glycosylation models and analysis tools.

Here we bring explicit attention to the concerns we raise above, we provide a focused, text-based representation of reaction rules that have been introduced for the purpose of formalizing these communications. GlycoCT^21^ and WURCS^22,23^ are two popular glycan nomenclatures in use today. GlycoCT was designed to maximize the descriptive specificity of the experimentally derived glycan structures data. WURCS, on the other hand, focuses on the uniqueness of a linear representation which promises efficient lookup in database queries. Both GlycoCT and WURCS produce unambiguous representations and are thereby invaluable for many applications, ranging from systems biology analyses^17^ to an international glycan structure repository^24–27^. GlycoCT and WURCS provide a high degree of unambiguous detail; however, they are limited in their human-readability. The glycan extension to IUPAC, on the other hand, is more human-readable^28^. It specifies the linkage and branch information in an intuitive and linear manner. In the hopes of mitigating the inconsistent application of IUPAC and inconvenient illustrations, Linear Code described a simplified version of IUPAC nomenclature^29^. Specifically, Linear Code is a syntax for representing glycoconjugates and their associated molecules in a simple linear fashion. While keeping the linkage and branch information, Linear Code removes the hyphens between monosaccharides and abbreviates the glycan symbols, thereby simplifying the representation without limiting flexibility. Given its readability and parsability, Linear Code has become a popular choice for representing reaction rules in computational models of glycan synthesis (Table 1). However, with the rise of Linear Code adaptations to represent reaction rules, we have seen increasing diversity in the syntax, including branch constraints, duplicate monosaccharides omission, logical operators, etc.

**Table 1.**
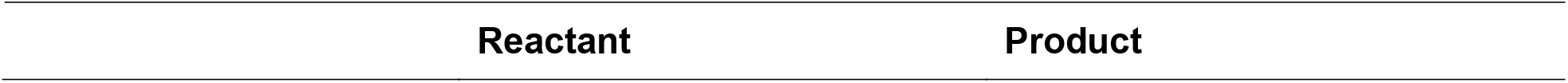
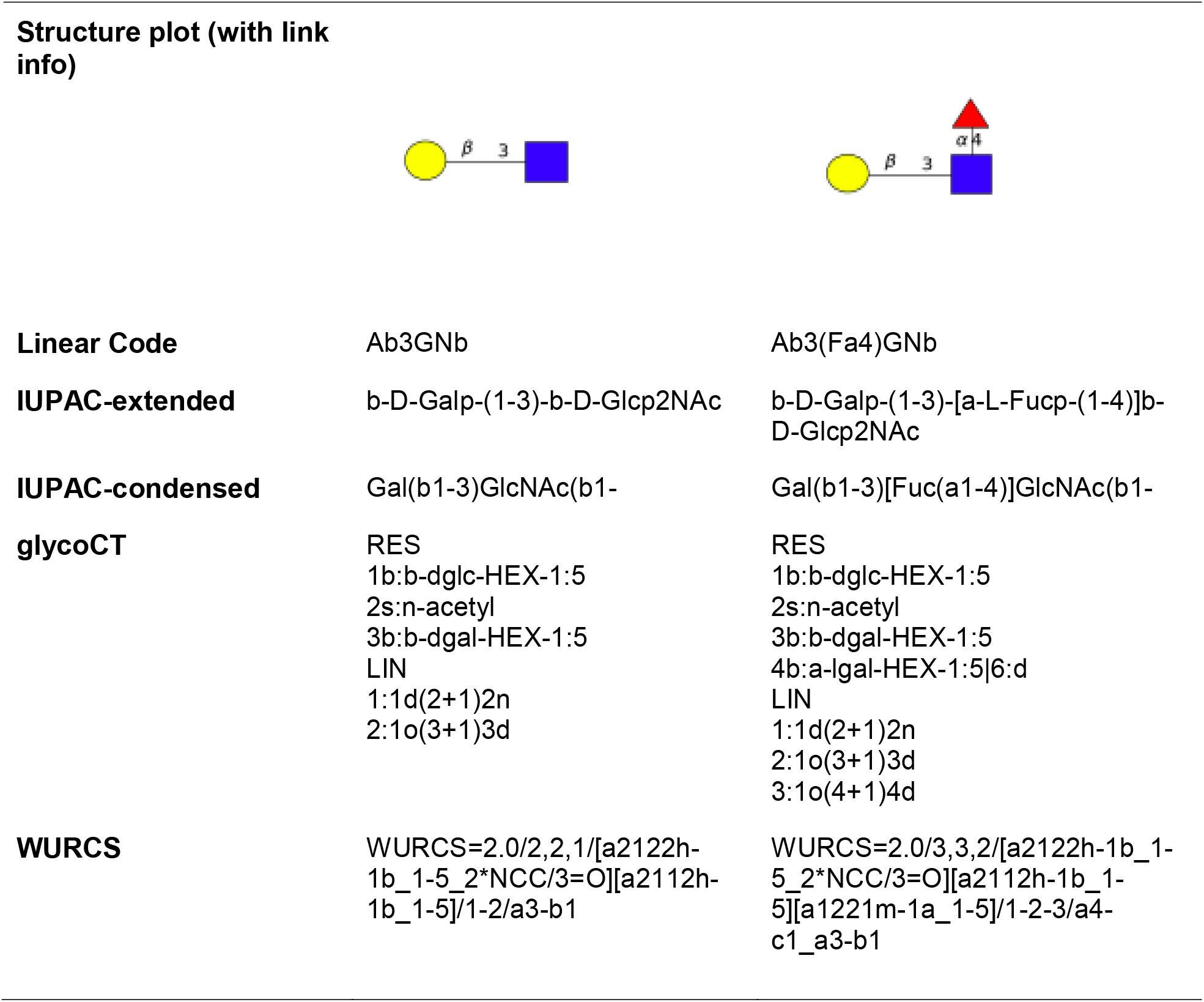
The reaction rule Ab3GNb → Ab3(Fa4)GNb represented in graph, Linear Code, IUPAC, GlycoCT and WURCS separately. Linear Code provides the most straightforward and succinct representation.

Here we critically review reaction rule nomenclature. In doing so, we seek to promote the development of a standardized and more unambiguous, readable and computable reaction rule representation. First, we examine the original usage of Linear Code for reaction rule representation by discussing six major categories of syntax rules. Second, we discuss the various adaptations that have been introduced in the current usage of Linear Code to represent reaction rules. Third, we further discuss the apparent nomenclature ambiguity emerging in the adaptation of Linear Code to systems glycobiology. Finally, we demonstrate the depth of the nomenclature crisis through the minimal overlap in presumably similar networks. While many solutions to this nomenclature might be offered, we focus on six major recommendations to provide a unified representation of reaction rules that are likely to have a broad impact on minimizing change to the current adaptations.

Common lore at universities describes architects who, rather than “prescribe” ideal paths for students through the mall, waited to see where students would walk. They built their paths over the trampled grass of the “descriptive” paths chosen by the students. Similarly, we intend to extend the thoughtful “prescription” of Linear Code to “descriptive” extensions that will comfortably accommodate those currently working in systems glycobiology. We also provide some key definitions for ease of reading (Figure 1, Table 2).

**Table 2.**
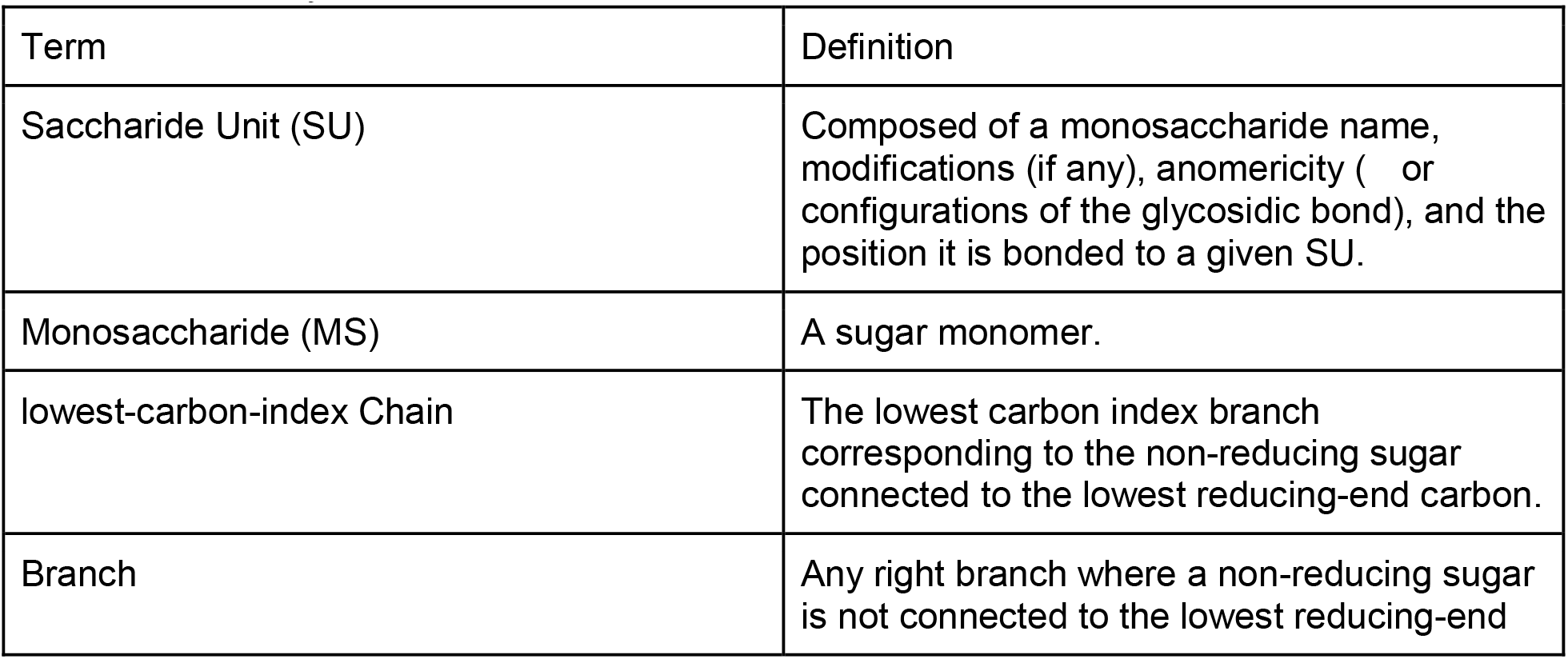
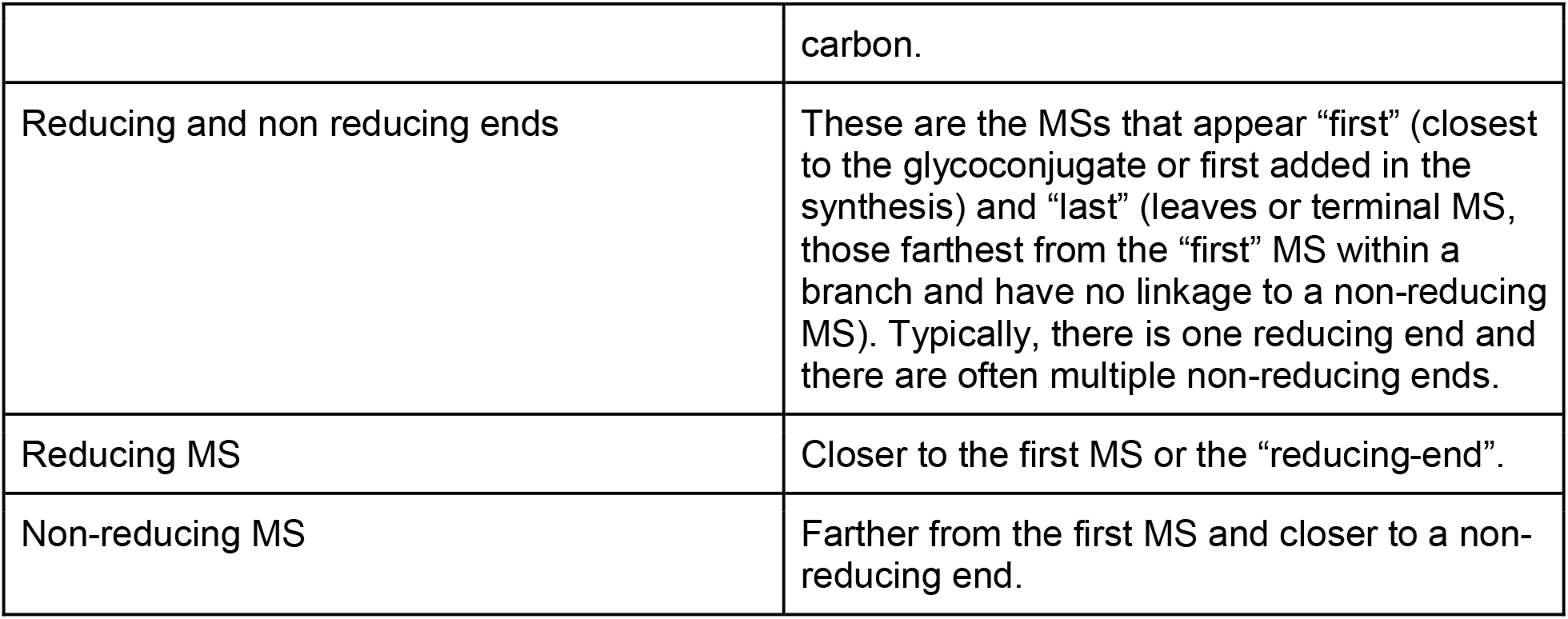
Glossary of essential terms

**Figure 1.**
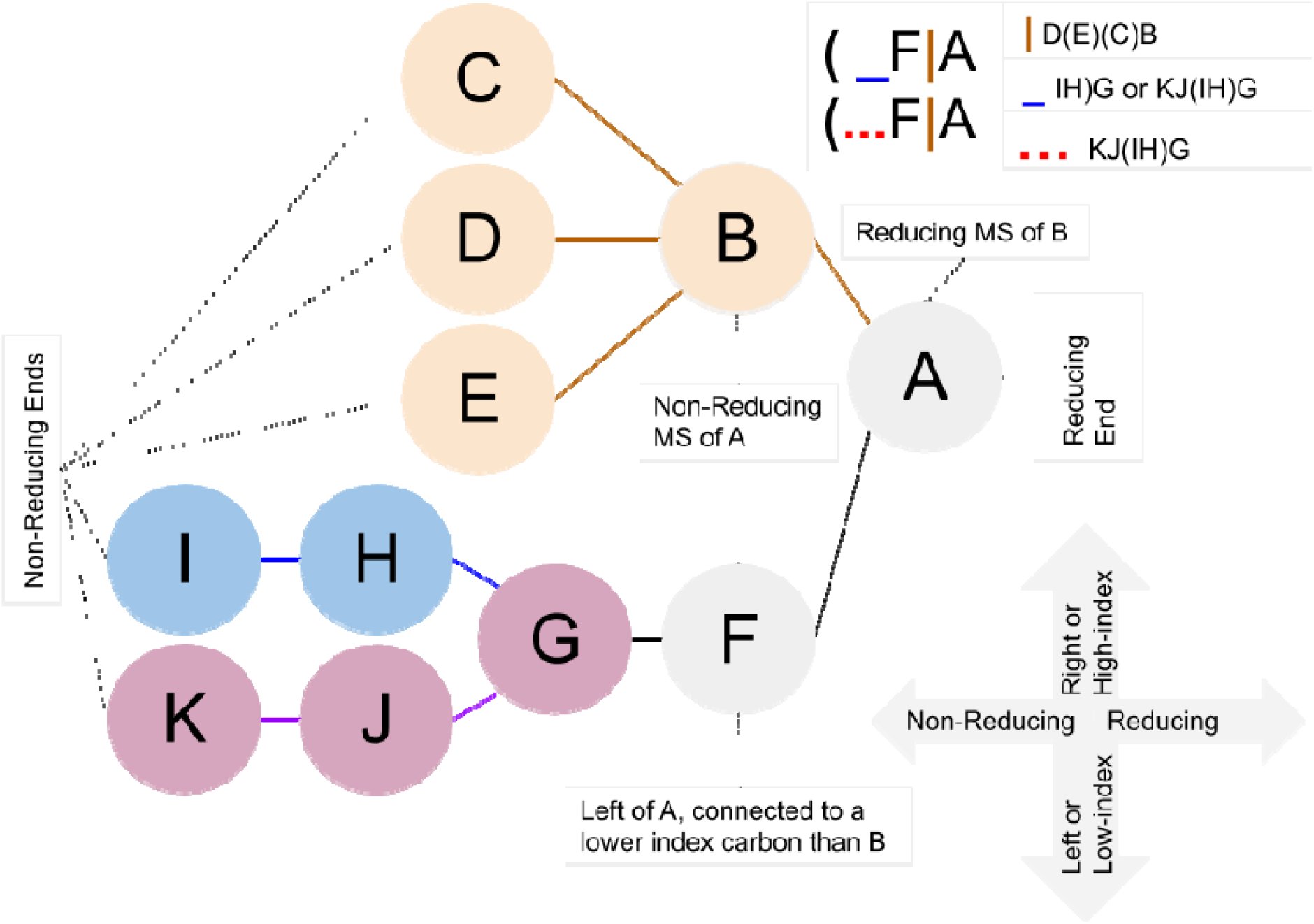
Common terminology and anatomy of a theoretical glycan, (KJ(IH)GF(D(E)(C)B)A. In this figure we demonstrate some key terminology as well as the three primary uncertainty operators: branch (orange), continuation (blue), and ligand (red). The structures matching these terms are shown in matching colors, those matching both the continuation and ligand are shown in purple. A ligand can typically be removed with one cut. A continuation is a connection from a node to a root that can “continue” or “bypasses” other branch points. The paths from I to G or K to G represent one continuation; to represent both paths, a continuation is necessary because traversing from I to G requires the syntactic “bypass” of the KJ branch.

## Syntax Rules of The Original Linear Code

Linear Code rules can be separated into six categories of syntax rules (Table 3): Stereospecificity & Ring structure rules (SRS), Modification rules (MR), Branch rules (BR), Repetition rules (RR), Glycoconjugate rules (GR), and Uncertainty rules (UR). The saccharide unit (SU) refers to a structure with four elements: anomericity, position number, modifications, and monosaccharide (MS).

○ ***Stereospecificity & Ring structure rules*** are set to differentiate the stereoisomers or distinct ring structures. A change from primary to secondary stereospecificity is denoted by “ ‘ ”, while a change to secondary ring structure is denoted “ ^ ”. A change to both secondary ring and stereospecificity is denoted “ ~ ”. For example, “ G ” represents glucopyranose, the pyranose conformation of glucose, with D stereospecificity. Glucopyranose with L stereospecificity is written as “ G’ ” (SRS1). Glucofuranose with D stereospecificity is written as “ G’ ” (SRS2), and glucofuranose with L specificity is written as “ G~ ” (SRS3). Similarly, galactofuranose, a common fungal monosaccharide, would be written “A^”
○ **Open Form rule** indicates that if the MS at the reducing end is open--a linear rather than cyclic MS, then the final character to the right of the string should be “o”. For example, lactose, galactose β-linked to glucose would be written as AbG if the reducing end glucose is closed and AbGo if the glucose is open; the open “o” takes the place of the linkage in this context. If the glucose is phosphorylated, this structure would be written AbG[P]o.
○ ***Modification rules*** specify a modification of MS at certain positions (MR1). MS + “ [” + modification + “] ” is used to denote the modification. For example, “G[2S]” describes sulfation on the second carbon of a D-glucopyranose. The anomericity is expressed to the right of the modification (i.e. “G[2S]a”). Multiple modifications to the same MS are ordered based on the position number inside the same brackets; ascending order from left to right. For some common modifications like *N*-acetylgalactosamine, instead of “A[2N],” Linear Code uses “AN” directly. Table 4 includes syntaxes of MS in Linear Code and common modified MSs. Common modification names can be found in Table 5.
○ ***Branch rules*** specify which non-reducing saccharide unit (SU) should be in the branch and which SU should continue the lowest-carbon-index chain; branching is determined by the identity of the first MS in a chain. When the non-reducing MSs are identical, the MS and its substituent chain, linked to the higher carbon of the reducing MS, will branch while the MS and substituent chain, linked to the lower carbon position of the same reducing MS, remains in the lowest-carbon-index chain (BR1). Otherwise, if the non-reducing MSs are different, the chain with a less frequent non-reducing MS (lower rank in Table 4) is considered the branch (BR2). The MS frequency is specified in Table 4, decreasing from top to bottom. When there are more than two non-reducing MSs linked to the same reducing MS, they are ranked, first by frequency, then by linkage index. The highest frequency MS is ranked higher, further to the left when the expression is written. Any MSs with equal rank after the frequency rank--those that are the same MS--are ranked by their linkage index, the lowest linkage indexes are ranked higher. A higher rank means these MSs, and their associated chains, will remain on the lowest-carbon-index chain, while the lower rank MSs will branch.
○ ***Repetition rules*** specify the contraction syntax for succinctly describing repeating MS units. The repetition structure is denoted by curly brackets, with a prefix of repetition times inside the brackets. For example, cellulose, which is a polymer of D-Glucose residues joined by b-1,4 linkages, is represented as “{nGb4}” (RR1). If a ring structure is repeated and the repeating unit is not connected “head to tail,” the MS where the repeating units are connected is marked between 2 dashes “ -- ” (RR2). An example is {nGa6Ga4(-Ab3-)Ub2Ha3Ha3Ha3}. Additionally, Banin et al. specifies that a cyclic motif, a form of repetition, is expressed using the letter “c”. While specification was limited in the original publication, we interpret “c” as denoting the “tail.” (-X-) denotes the head if it is not the left end and “c” denotes the tail if it is not the right end of the string. For example, in the molecule nGa6Ga4(-Ab3-)Ub2Ha3Ha3Ha3, Ab3 connects to the reducing end, Ha3. But if Ab3 was connected to the second Ha3 from right instead, we can specify the point of the cycle using a “c,” nGa6Ga4(-Ab3-)Ub2Ha3H<u>**c**</u>a3Ha3.
○ ***Glycoconjugate rules*** describe when a reducing end of a SU is connected to non-carbohydrate moieties, Glycoconjugate rules regulate that amino acid sequences are written after “ ; ”, lipid moieties are written after “ : ”, and other glycosides are written after “ # ” (GR1). For example, a glucose β-linked to a Ceramide is written as “Gb:C.”
○ ***Uncertainty rules*** describe syntax for when certain features of the SU are unknown or have more than one possibility. If the anomericity of certain bonds is unknown, Linear Code uses “ ? ” (i.e. AN?3G) (UR1). If both linkage anomericity and position are unknown, Linear Code uses “ ?? ” (i.e. AN??G) (UR2). If an entire SU is unknown, “ * ” can be used instead. ANb3*A represents a three SU glycan, where the second SU is unknown (UR3). When two monosaccharides are possible for a given SU, Linear Code uses the forward slash to separate them. When SU ambiguity refers to anomericity, position number, modifications, or MS, a single “ / ” is used (i.e. ANb3/4) (UR4). Given two complete possible SUs, Linear Code uses “ // ” to separate them (i.e. Ab4//Ga2Aa3 represents Ab4Aa3 or Ga2Aa3) (UR5). When analyzing fragmented glycans, an “< index number>%” is used to store fragmented structures as a variable. For example, NNa6=1%|1%Ab4GNb2Ma3(1%Ab4GNb2Ma6)Mb4Gb is a glycan containing a terminal α2,6-linked sialic acid (NNa6) whose linkage position is unknown. Here, the “ | ” is used to separate the fragment(s) and core structure components (UR6).

**Table 3.**
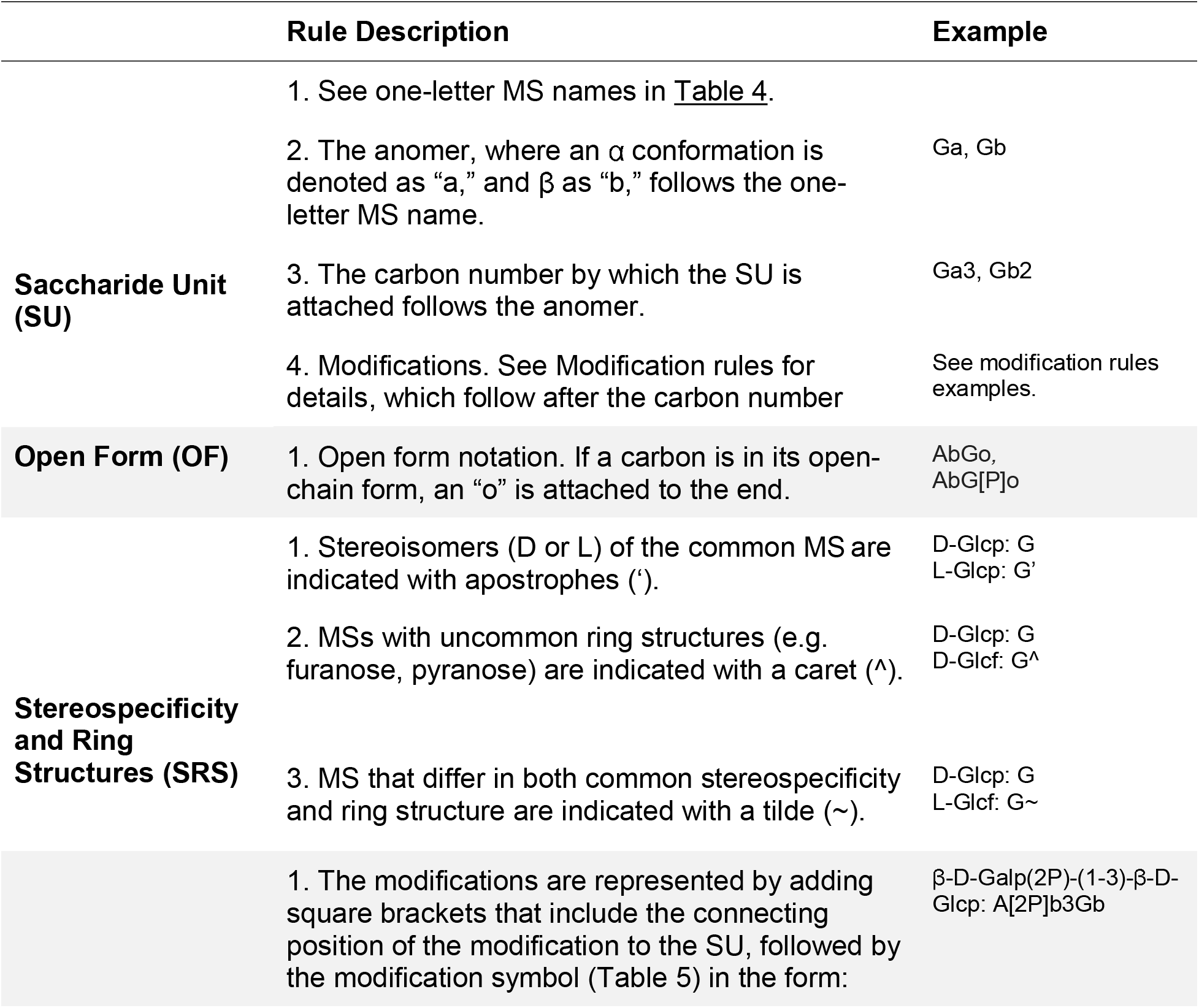
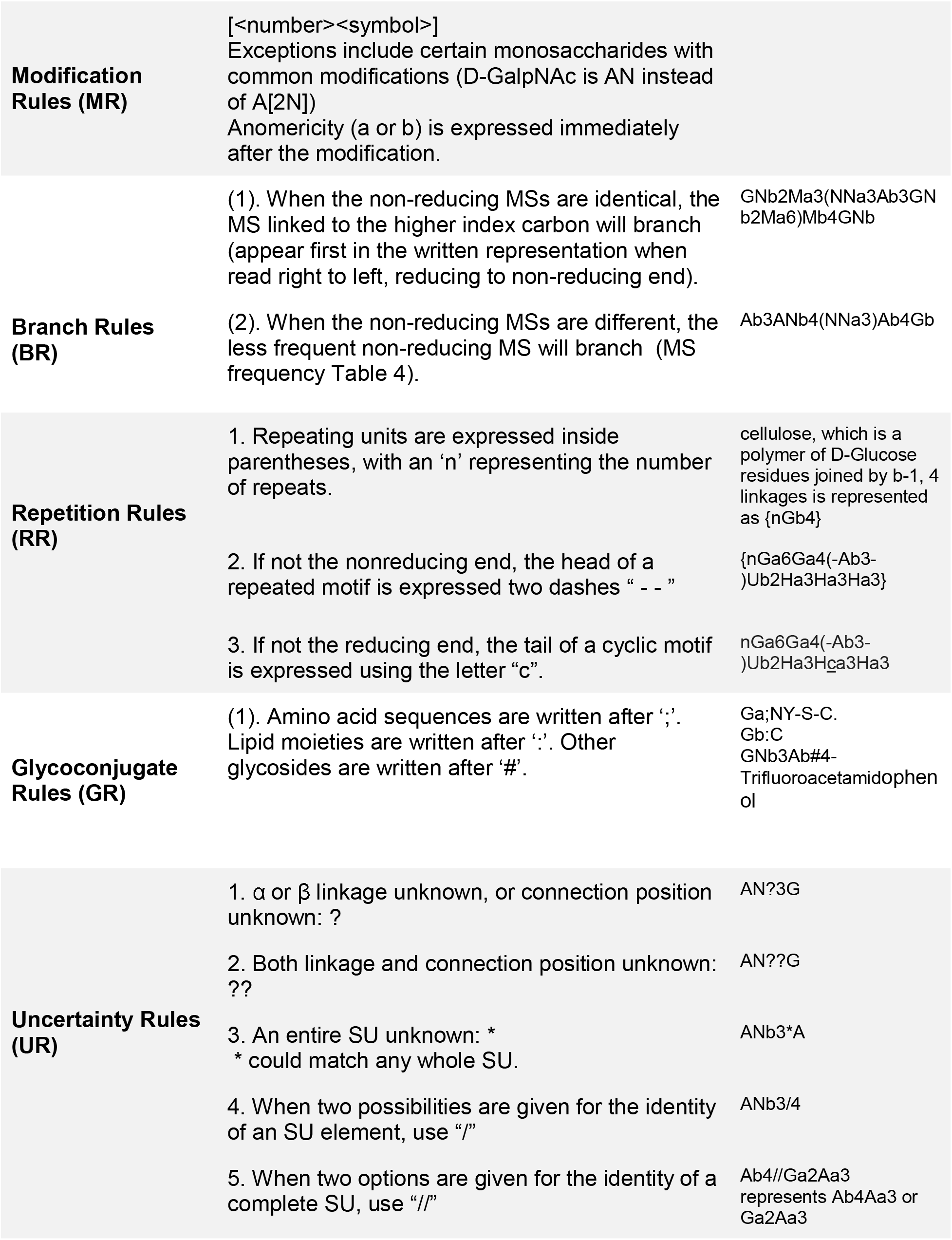
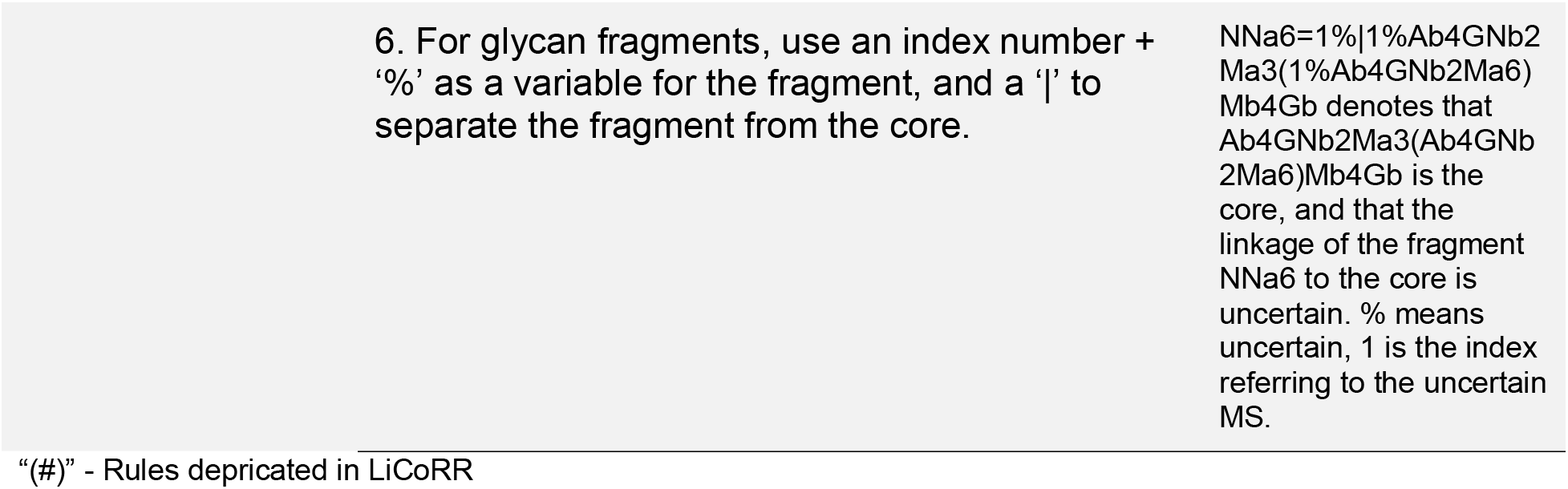
Original Linear Code rules^29^

**Table 4.** Common monosaccharides and their Linear Codes (adapted from^29^). We have added NG as it has become a clearly important monosaccharide excluded from the original list. Full monosaccharide descriptions are recorded in IUPAC^18^; all terms can be found at https://www.qmul.ac.uk/sbcs/iupac/2carb/38.html

| Monosaccharides<sup>1</sup> | Linear Code |
| --- | --- |
| D-Glucose | G |
| D-Galactose | A |
| N-Acetylglucosamine | GN |
| N-Acetylgalactosamine | AN |
| D-Mannose | M |
| N-Acetylneuraminic acid | NN |
| *N-Glycolylneuraminic acid | NG |
| Neuraminic acid | N |
| 2-Keto-3-deoxynononic acid | K |
| 3-deoxy-D-manno-2 Octulopyranosylonic acid | W |
| D-Galacturonic acid | L |
| D-Iduronic acid | I |
| L-Rhamnose | H |
| L-Fucose | F |
| D-Xylose | X |
| D-Ribose | B |
| L-Arabinofuranose | R |
| D-Glucuronic acid | U |
| D-Allose | O |
| D-Apiose | P |
| D-Fructofuranose | E |
| *Ascarylose | C |
| *Ribitol | T |

1 - All the monosaccharides are in their pyranose form unless otherwise noted.
2 - Asterisk ("\*") represents an update from the original table.

**Table 5.** Common modifications and their Linear Code (from^29^).

| Modification Type | Linear Code |
| --- | --- |
| Deacetylated N-acetyl | Q |
| Phosphoethanolamine | PE |
| Inositol | IN |
| Methyl | ME |
| N-acetyl | N |
| O-acetyl | T |
| Phosphate | P |
| Phosphocholine | PC |
| Pyruvate | PYR |
| Sulfate | S |
| Sulfide | SH |
| Aminoethylphosphonate | EP |
| *Deoxy | D |
| *Carboxylic Acid | CA |
| *Amine | A |
| *Amide | AO |
| *Ketone | K |

1 - Asterisk ("\*") represents an update from the original table.

## Current Usage of Linear Code to represent Reaction Rules

Linear Code was first used to represent reaction rules in 2009. A reaction network, specifying glycans with condensed IUPAC and Linear Code, was trained on mass spectrometry abundance to learn biosynthetic enzyme activities^10^. Their reaction rules table contained four features: enzyme, reactant, product, and constraint. For the implementation rules, not all original Linear Code rules are adopted. Krambeck et al. maintained the linkage information (Table 3: SU2), one-letter MS abbreviation (Table 4), and branch rules (Table 3: BR), which are the necessary conditions to denote a glycan with branches. On the other hand, symbols “ ~ ”, “ * ”, “ | ” were defined with new meanings, though they already had their meanings in the original Linear Code rules (Table 3: SRS3, UR3, UR6 respectively). Instead, Krambeck et al. introduced several new symbols to convey logical relationships (“ & ”, “ ~ ”, “or”) and structural ambiguity (“ … ”, “ _ ”, “ | ”, “ * ”), all of which were used to specify constraints. For example, a constraint “Ma6 & Ma3” means the reaction will happen only if both Ma6 and Ma3 appear in the glycan; as an N-glycans, these are the terminal mannoses capping the chitobiose core. The “Ma6 or Ma3” constraint promotes the reaction if either Ma6 or Ma3 exists. “~Ma6” means the reaction will not happen if Ma6 is present in the glycan. The structure denotations are indicators of certain parts of the glycan. “ … ” can be replaced with either nothing or any polysaccharide with matched parenthesis. “ _ ”, in Krambeck et al., can be replaced with either nothing or any polysaccharide where each left parenthesis is matched to a right parenthesis but where right parentheses are not necessarily matched. “ | ” represents a possible branch. We expand on the distinctions between “ … ”, “ _ ” and “ | ” in a later section “Substring uncertainty operators” (Table 4). The asterisk “ * ” stands for the reaction site, which is the position where the new MS will be added or an MS is removed. Krambeck et al. also uses “ # ” to describe constraints around the number of MS that may appear in a glycan. For example, the constraint “#A = 0” means the reaction will happen only if there is no galactose. The Krambeck et al. adaptation is the most common adaptation of Linear Code to represent reaction rules^7,13,15,30^.

Based on the Linear Code reaction rules framework Krambeck et al. created, later researchers introduced new attributions that specify and simplify the description of reactions. Bennun et al. and Spahn et al. include the amino acid at the end of the Linear Code attached by a semicolon “ ; ”. This suffix is exactly the syntax from the original Linear Code rules (Table 3: GR1). The reaction rules table generated by Spahn et al. also provided localization information, which is either *cis*, *trans* or *medial* to denote the Golgi compartment where the reactions happen^13,14^. The subcellular localization of a reaction, in the endoplasmic reticulum, golgi, cytoplasm (bacteria and archaea) or lysosome (degradation, Man-6-P dephosphorylation and lysosomal glycoprotein biosynthesis ^31,32^ or paucimannose recycling^33^), is an important constraints on glycosylation^34^, therefore, the addition of this information to the Linear Code reaction rules provides insights into the glycosylation types.

Some models of glycan synthesis generated reaction rule tables with an additional column Enzyme Commission number (EC number)^7,16,35^. The EC number system is a numerical classification scheme for enzyme-catalyzed reactions that provides an unambiguous accession to a cataloged reaction^36^. The inclusion of an EC number in the reaction rules table, therefore, promotes the clarity, interoperability, and reproducibility of the generated reaction model.

A common syntax used by most studies is the leftmost “ (” to represent the terminal, non-reducing end of the glycan chain. It specifies whether the leftmost MS is the terminal MS both visually and computationally. For example, the reaction rule (GN → (Ab3GN applies to all reactions which add one galactose to a terminal *N*-acetylglucosamine. On the other hand, the reaction rule GN → Ab3GN applies to all reactions which add a galactose to an *N*-acetylglucosamine, but not necessarily the terminal one. The leftmost “ (”, therefore, can easily vary the glycan substrate substantially.

Though Linear Code was developed with parsability in mind, some have found it useful to make a specific computational implementation of the reaction rules to accommodate the syntactic constraint of programming languages. A human milk oligosaccharide metaglycome was constructed using a combination of linear code, glycan structures represented in XML and XPath queries^37^. Separately, Akune et al. generated a theoretical *N*-glycan database called UniCorn, based upon a Perl implementation of reactions on glycans represented in Linear Code^35^. Though Linear Code is computer-parsable, there is still substantial work necessary to implement that parsing because there is no standard representation for handling the wide variety of reactions possible, nor open-source software available to implement the parsing of such rules.

Representing reaction rules in Linear Code is not easy because of a few ambiguous cases not completely described in the initial Linear Code paper. Subsequent studies, therefore, have developed their own ways to idealize reaction rule implementations based on Linear Code. Using the framework Krambeck et al. built^10^, new information like Golgi localization and EC numbers are added to specify and simplify the reaction rules.

## Original Prescriptions for Substring Uncertainty Operators

In its original conception ^10^, the adaptation of Linear Code to represent reaction rules aimed to describe how glycosylation enzymes change the structure of glycans in terms of how the Linear Code character string descriptions of the glycans are changed (**Figure 1**). In the simplest case, we can specify a substring of the substrate code to be replaced by a new substring to form the product code. In addition, there can be constraint and adjustment substrings whose presence or absence within the substrate string either restricts which glycans can be substrates of a particular enzyme or modifies the reaction rate parameters. Uncertainty operators have been developed to facilitate searching substrings for specific structural features of a glycan implied by the substrings.

The substring specifications for the substrate, product and adjustments can include any combination of characters included in the glycan codes in addition to uncertainty operators inserted within the directly specified characters. Each uncertainty operator is represented by one or more characters, such as “…” or “_” (Table 6). To perform substring matching of a glycan to a substring with uncertainty operators, we first identify the characters of the specified string immediately before and after the uncertainty operator. If found, we then test the substring of the glycan string between these two matched character strings and check for the appropriate uncertainty operator properties. In parsing the glycan code, an initial left parenthesis is always added to the complete glycan code so that the terminal end of every branch of the glycan is always a left parenthesis. Below, in defining the properties of substrings corresponding to an uncertainty operator, we use the symbol X to represent some monosaccharide with its connection, such as Ma3, GNb4, etc.

**Table 6.**
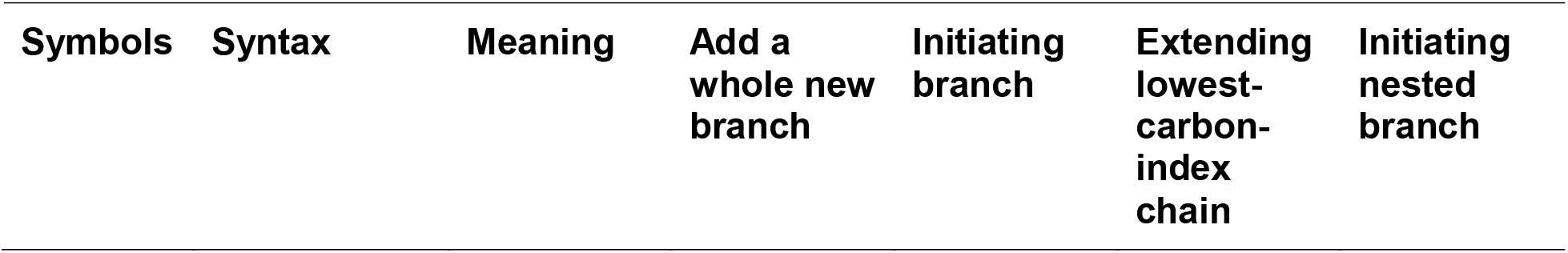
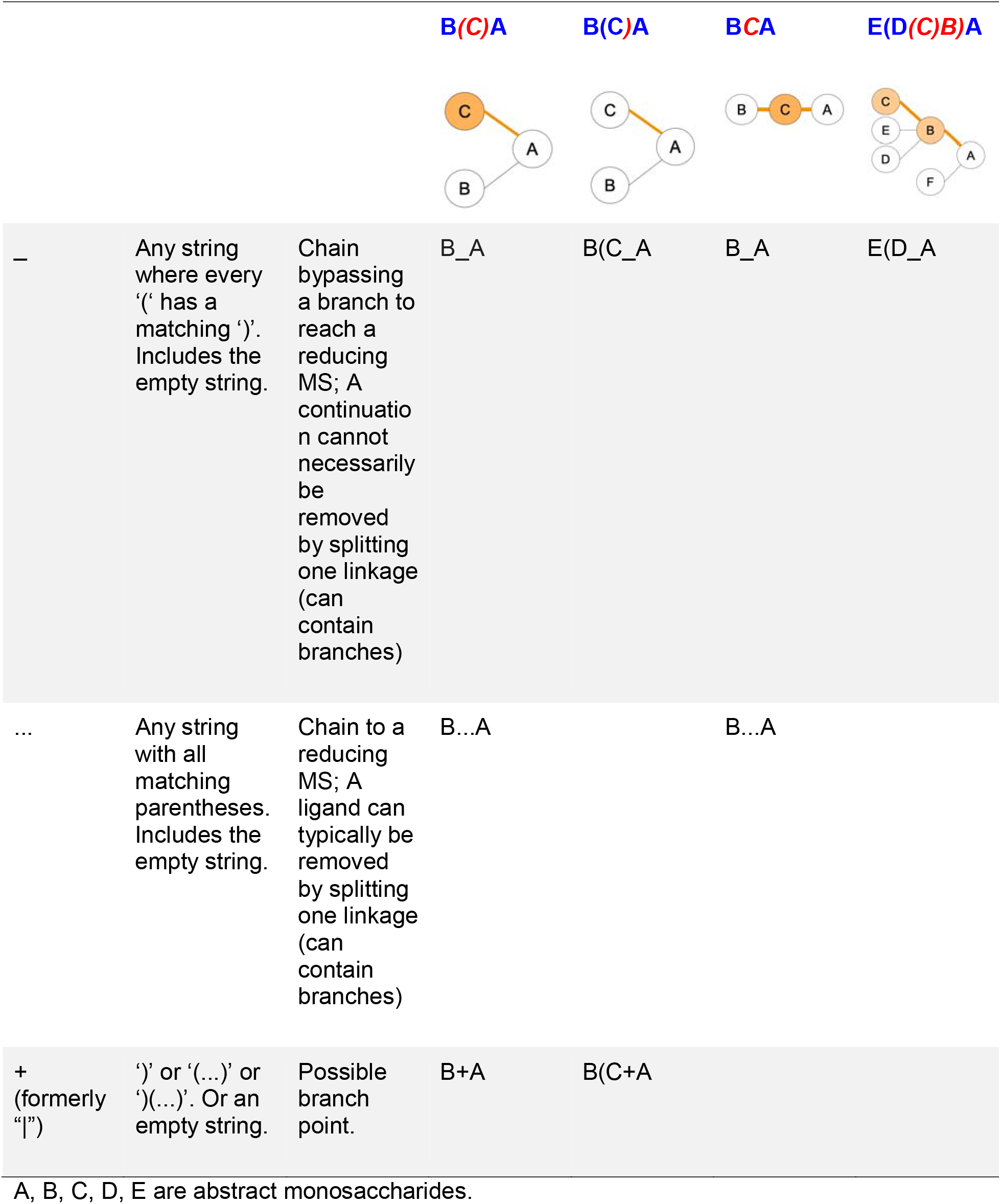
The difference between “ _ ”, “ … ” and “ | ” with illustrations. These symbols were proposed by Krambeck et al^10^. The initial names are ligand (“ … ”), continuation (“ _ ”), and possible branch (“ | ”). Each uncertainty operator in the last four example columns can be replaced by the substring in red to achieve the behavior described in the column header. For a more comprehensive look at the usage of these uncertainty operators, see Supplemental Table S1 for a manual collection of matches, and Table S4 for an automated collection of matches.

There are three types of uncertainty operators. The ligand “ …”, the continuation “_”, and the branch “ | ”. Each has a specific syntactic match, but intuitively, the ligand is a chain that can contain branches, the continuation is a chain that can include branches terminated outside of the continuation, and the branch is either a complete branch or nothing. Functionally, “…” indicates the “leftward” extension along the lowest-carbon-index chain, “|” indicates the “rightward” extension along the highest-carbon-index chain, and “_” indicates an extension along either the left for right chain (Figure 1).

More specifically, (1) the ligand uncertainty operator indicates a chain of MSs that can include attached branches completely contained in the substring, (2) the continuation uncertainty operator indicates a chain of MSs that can include attached branches that may not be wholly contained in the substring, and (3) the possible branch uncertainty operator indicates where a branch may be included in the substring. Due to the nuances of representing a glycan linearly, these are not complete definitions.

**Ligand “…”** - A ligand is a fragment of a larger molecule connected to the rest of the molecule at one point. Glycans are themselves ligands, as they are pieces of larger molecules. A substring is a valid “ligand” if each parenthesis in that substring is uniquely and appropriately matched; each left parenthesis must be followed by a corresponding right parenthesis and each right parenthesis must be preceded by a corresponding left parenthesis: “)(“ are not matched parentheses. Any substring with all left and right parentheses matched, including an empty string, is considered a ligand. If we select a substring of the code representing a glycan it may or may not represent a ligand. For example, XXX)X, XX(XX, X)(XX are not valid ligands, while XX(XXX)XX is valid.

Functionally, ligands can serve as connectors between the left and right portions of a glycan a user would like to specify. A ligand is simply a chain of monosaccharides which may contain nested branches; the nested branches must also be ligands. However, there can be many chains or paths through a ligand, starting from one of the terminal monosaccharides and culminating at the root end; there are many ligands within most ligands. Any of these ligands can serve as a connector from the root (reducing) end of one ligand to a terminal (non-reducing) end of another. The key property of a ligand substring is that all the included branches of the ligand are completely contained in the substring.

**Continuation “_”-** As we parse from left to right through a substring, we may find left parenthesis (entering into a branch) and right parenthesis (exiting a branch). A ligand, with matched parentheses, indicates an equal number of branch initiations and completions. On the other hand, a substring with an unmatched right parenthesis, for example, XXX)XX or X)(XX)XX, indicates a net termination of branching; each right parenthesis indicates moving out of a branch towards a root. As long as all the left parentheses encountered are followed by right parentheses, we are following a path along a connected chain of the glycan structure. A substring where every left parenthesis can be matched with a following right parenthesis, but not necessarily vice versa, is a “continuation.” Again, we include the empty string in this class of substrings. Note that any ligand is also a continuation.

The continuation uncertainty operator can be very useful in formulating rules that apply to specific monosaccharides connected by a chain of monosaccharides to a particular reducing monosaccharide of the glycan structure. For example, the iGnT enzyme adds a Gnb3 group to a terminal galactose group and has a preference for the two branches that are connected to the Ma6 of the root Ma3(Ma6)Mb4 structure. This leads to an adjustment rule based on the string Ma3|(*_Ma6)Mb4. Here the “|” uncertainty operator is used to allow for the possible presence of a bisecting GlcNAc on the root mannose: Ma3(GNb4)(…Ma6)Mb4. The “*” indicates the site of the enzyme action.

**Possible branch “|” -** As discussed, parsing a linear glycan from left to right, we can encounter matched parentheses indicative of a ligand or unmatched right parentheses indicative of a closing branch. We can leverage the branch closer offered by these symbols to mandate a possible branch. The definition of the “possible branch” is one of either: “) ”, “ (…) ”, “)(…) ” or an empty string. This uncertainty operator can be replaced with either a branch, the start of a branch or nothing. It allows the same specification string to work whether an additional branch is present at the position of the uncertainty operator or not, as in the above example.

## Divergence in Current Implementations of Reaction Rules from Original Linear Code

Linear Code is a useful notation to succinctly describe glycan structures. It is thus powerful to represent reactants, products and reaction constraints. Constraint syntax (e.g., the glycan site where a reaction happens, and logical relationships), however, was beyond the scope of the original Linear Code rules. Therefore, different adaptations are introduced throughout the literature.

We have identified four symbols that are prescribed with different meanings than they were originally assigned in the Linear Code rules:

○ ***Ambiguous symbol 1*** – Originally, “ ~ ” following an MS name was used to denote the MS with different stereospecificity and ring structure from the common form (Table 3: SRS3). For example, while “ G ” represents D-glucopyranose, “ G~ ” represents L-glucofuranose, which is rarely seen. To represent reaction rules, “ ~ ” was used, instead, to convey logical negation^7,10,13,30^. For example, a constraint “~Ab” means the reaction will not happen if a β-galactose is present.
○ ***Ambiguous symbol 2*** – “ | ”. For reaction rules, “ | ” is widely used to represent a potential branched structure in a substrate^7,10,13,30^. For example, “(GNb2|Ma3” represents a glycan structure with a potential branch on the mannose. However, “ | ” was originally designed to separate the certain and uncertain parts in a fragmented glycan, where there is a possibility of different structures (UR6).
○ ***Ambiguous symbol 3*** – “ # ”. Originally, “ # ” was designated to signify the starting point of glycosides that are not amino acids or lipid moieties (Table 3: GR1). For example, “GNb3Ab” connected to a “4-Trifluoroacetamidophenol” is written as “GNb3Ab#4-Trifluoroacetamidophenol.”
○ ***Ambiguous symbol 4*** – “ * ”. Another ambiguous symbol is the asterisk “ * ”. In the original Linear Code context, “ * ” is used when an entire saccharide unit in the complex carbohydrate is unknown. In reaction rules representation, “ * ” marks the “reaction site”, the position of the first difference between product and substrate strings in Linear Code form^7,10,13,30^. Note that the “reaction site” does not necessarily refer to the exact place that the reaction happens. For example, given the reaction “(…Ab4GNb → (Fa3(…Ab4)GNb,” the constraint “ (*Ab4 or (*Fa2Ab4” means that the reaction will happen if and only if the “ … ” in the reactant represents either nothing or “Fa2.” In this case, “ * ” on the left of “Ab4” indicates where the reactant and the product differ from left to right in the Linear Code expression. However, the real reaction takes place at the “GNb,” not “Ab4.” Demonstrating the left-to-right specificity of “ * ”, consider the rule, (Ma2Ma → (Ma with constraint ~*2Ma3(…Ma6)Ma6. This constraint rules out removing the Ma2 on the middle branch of the original M9 glycan, Ma2Ma2Ma3(Ma2Ma3(Ma2Ma6)Ma6)Mb4GNb4GN;Asn. If parsed from right to left, the constraint would be ~*Ma3(…Ma6)Ma6.

Linear Code is primarily a representation of glycan structure, and the formulation of reaction rules from Linear Code emerged as it was adapted for use with systems biology reaction networks. Specifically, when researchers aimed to define rules for reactions when building the networks, additional symbols were needed and, therefore, proposed. However, these now differ between studies.

In the first study, building reaction networks from Linear Code, Krambeck et al. defined “ … ”, “ _ ”, and “ | ” as uncertainty operators to indicate specific combinations or balanced or unbalanced (complete or incomplete) branches^7,10,30^. Spahn et al. used only two of the three symbols; “ | ” to indicate branching and “ … ” to represent continuation^13^. In this section, we will only focus on Krambeck et al. syntax. Syntactically, each of these symbols specifies whether or not the monosaccharides following the symbol, the first monosaccharide within the uncertainty operator replacement, appear within parentheses. If the monosaccharides appear within parentheses, it is “branching” off the lowest-carbon-index chain; otherwise, it is a “continuation” along the lowest-carbon-index chain. Each uncertainty operator describes a branching and/or continuation. Additionally, an uncertainty operator can require a complete phrase, with matched parentheses, or not. Finally, some uncertainty operators can be replaced with nothing (the empty string).

In the original Krambeck et al. implementation, multiple disjunctive constraints are connected by the logical disjunction “or.” An example is “(*Ab4 or (*NNa3Ab4” (Table 7). In the Liang et al. adaptation, however, the “or” relationship is delineated by writing each reaction rule on separate lines. For example, the two constraints for the reaction rule “(…Ab4GNb → (Fa3(…Ab4)GNb” would simply be written on two lines. (Table 8).

**Table 7.**
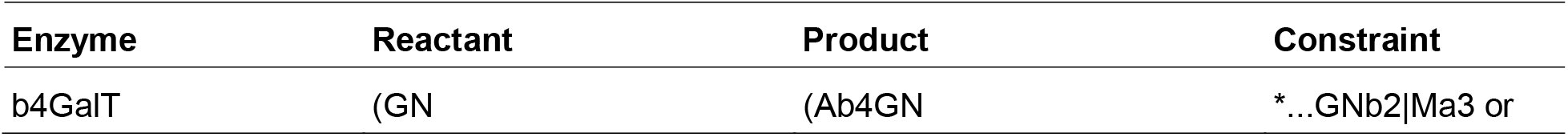
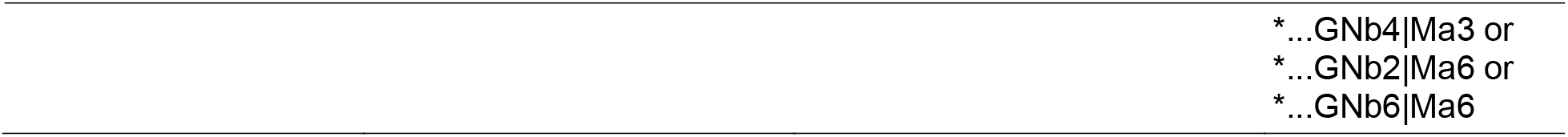
The reaction rule (GN → (Ab4GN with four constraints written in the same cell.

**Table 8.**
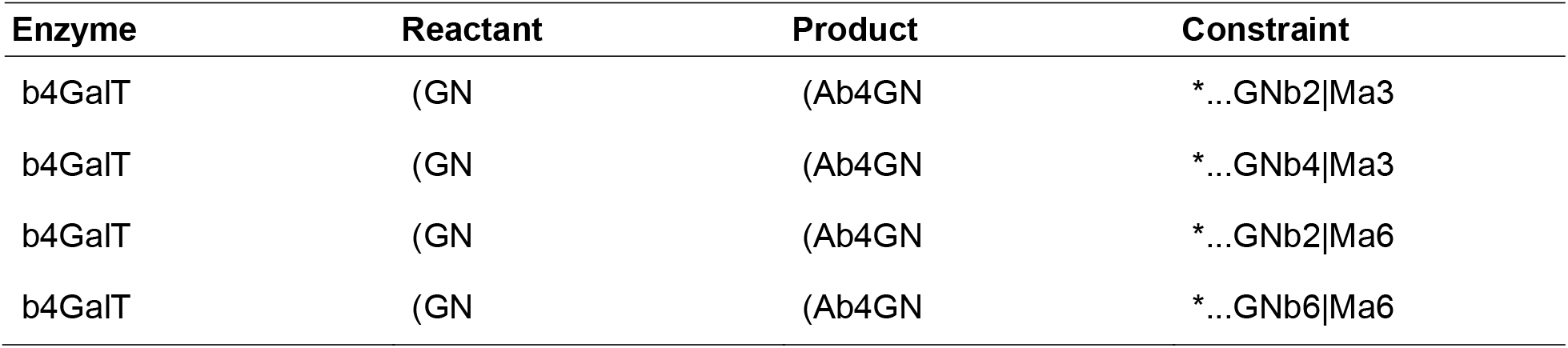
The reaction rule (GN → (Ab4GN with four constraints written on separate lines.

Most adaptations of reaction rule implementations are more or less related to the earliest Krambeck et al. adaptation. Some symbols are only seen in the Krambeck et al. adaptation. Besides the “ # ” as the number symbol, Krambeck et al. also uses “Gnbis” to refer to the specific structure of bisecting GN, which is “Ma3(GNb4)(…Ma6)Mb4.”

Several reaction rules for N-glycan biosynthesis are presented for direct comparison (Table 9, Table S5). While there were several apparent divergences in the usage of terms, the rules are predominantly similar. The intent of this paper is to ensure the consistency of these rulesets going forward.

**Table 9.**
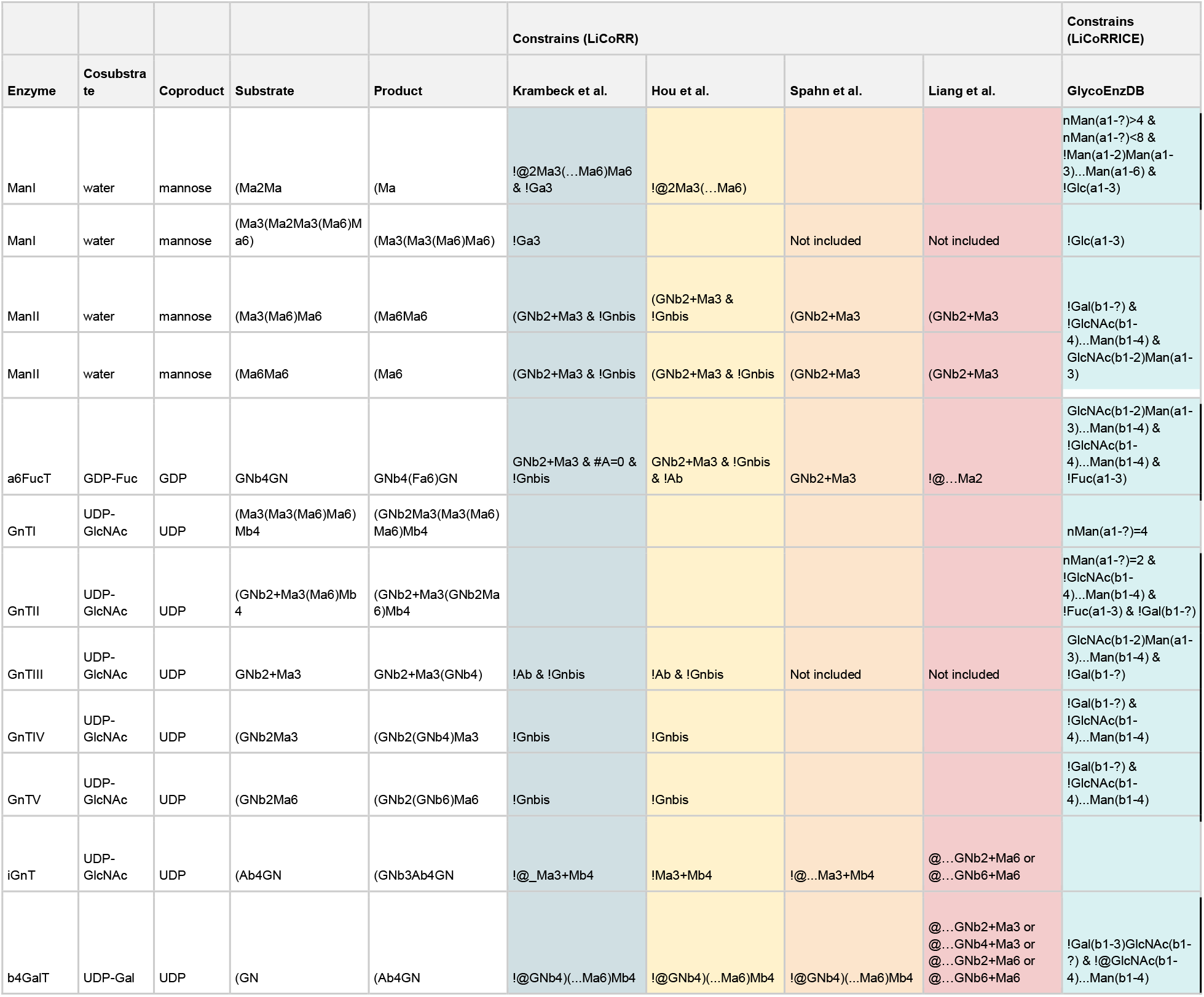
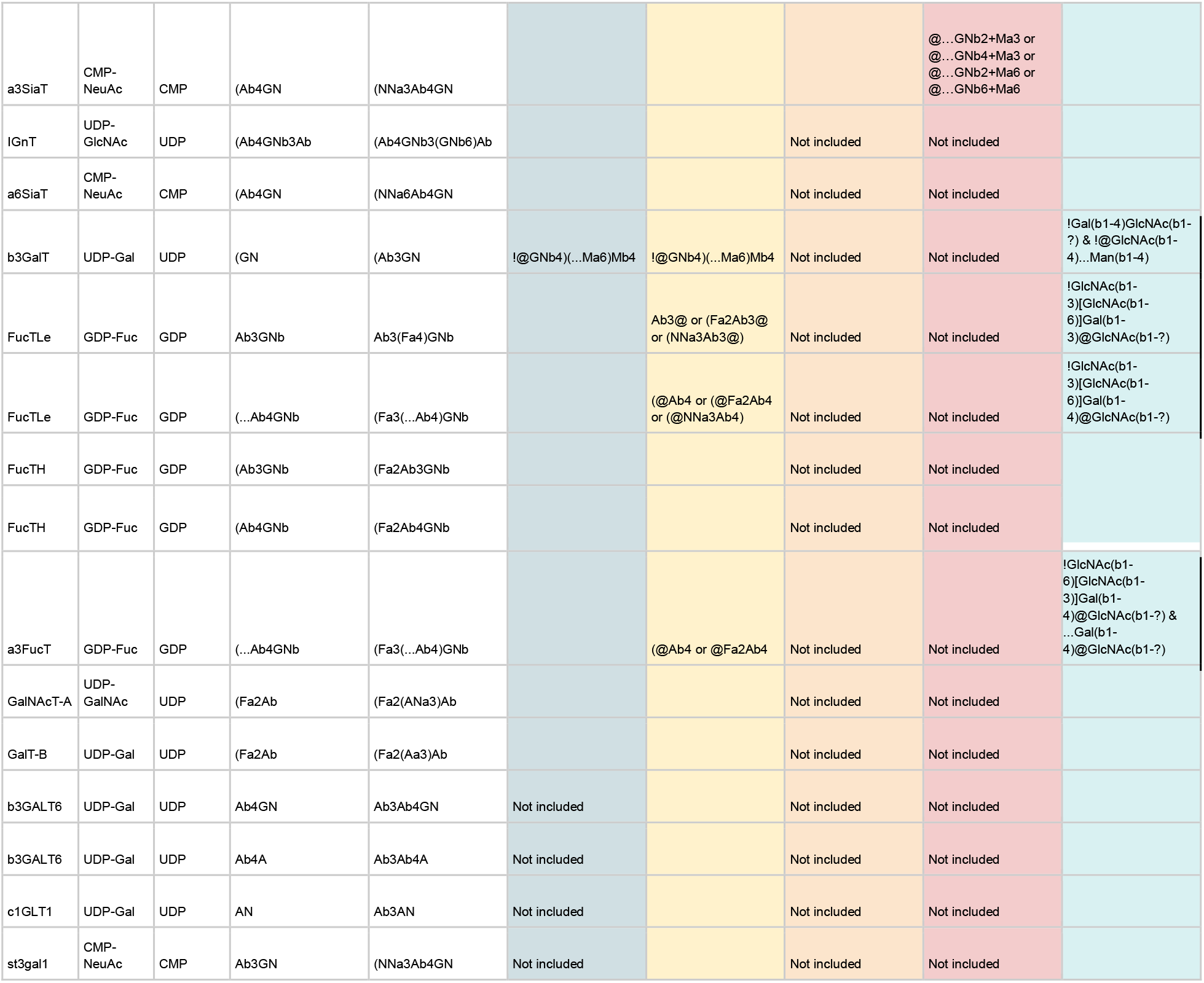
Reaction rules from multiple N-glycan biosynthesis models in LiCoRR representation. This table can be found in Linear Code in Table S5.

## Recommendations to Unify Descriptive Usages of Linear Code for Reaction Rules (*LiCoRR*)

Linear Code has shown its utility for the compact description of glycans and compatibility with efforts to define glycan reaction rules for systems biology models. A few ambiguities have emerged through different interpretations and implementations. Here we propose possible solutions as described by the original prescription for Linear Code, the consensus of the community, and our recommendation following this survey.

We have demonstrated the LiCoRR representation of all N-glycosylation reaction rules discussed in this paper in Table 9. Table 9 also includes an instance of these reaction rules written with IUPAC monosaccharides and linkages from GlycoEnzDB. Due to incomplete adoption and flexibility of Linear Code monosaccharides, we encourage users to accommodate both Linear Code and IUPAC monosaccharides when possible to facilitate interoperability; Linear Code monosaccharides may not be sufficient for every project while IUPAC-extended nomenclature^18^ is actively maintained to ensure complete coverage of known sugars. If a user wants to specify that they are using LiCoRR with IUPAC monosaccharides, they can specify it as “LiCoRRICE” the LiCoRR-IUPAC Complement Expression. We also provide the matched constraints in Table 9 as original Linear Code (Table S5). It should be noted that IUPAC uses square brackets, “[]”, rather than parentheses, “()”, to delineate branching. Therefore, the wildcards should recognize square brackets rather than parentheses. Additionally, IUPAC does not use deterministic branching. Therefore, specifying branch direction is not meaningful and the three branch-specific LiCoRR wildcards can be reduced to one, “…”, in LiCoRRICE. With these small changes, LiCoRR can be extended to LiCoRRICE and, as such, gain access to its carefully curated and growing list of MS units and modifications.

The original Linear Code syntax contains eighteen specific regulations across seven categories, among which only five regulations are seen in reaction rule implementations. In fact, the five regulations include three SU elements (MS name, linkage-type, position number), denotations (Table 3: SU) and one branch rule (Table 3: BR1). BR1 dictates that when two branching MSs are identical, the MS linked to the higher index carbon will have its chain on the branch (Table 3: BR1). If we extend the condition for BR1 from identical MSs to all MSs, written glycan structures will still maintain their uniqueness since each position on the MS can only connect to a single MS. BR2 dictates that the least frequent MS of the pair will branch (Table 3: BR2). BR2 solves the case when there are more than two non-reducing MSs linked to the same reducing MS. However, if we applied the expanded BR1 and ordered the chains based on decreasing position numbers from right to left in multi-chain cases, BR2 would be redundant. For example, Ab4(GNb4GNb3)(GNb6)Ab4Gb will be written as GNb4GNb3(Ab4)(GNb6)Ab4Gb.

Among the logical relationships required for constraint specification, only “or” is seen in the original Linear Code rules. “ / ” was designed to separate two possibilities within an SU (Table 3: UR4) and “ // ” was used to separate two possible complete SU options (Table 3: UR5). It would cause unnecessary confusion if “ / ” and “ // ” are used to denote the “or” relationship between constraints. Therefore, the task to convey Boolean logic among constraints was left to emerge organically in its application to reaction rules.

○ ***Recommendation 1*** – “*Logical negation*.” The field chose to use the “ ~ ” to indicate logical negation (Table 10: a). Unfortunately, this choice conflicts with the ability to express an uncommon stereospecificity, as prescribed in the original Linear Code (Table 3: SRS3). Though this is a rare necessity, and the original linear code tilde appears on the right of the monosaccharide, usage of a “ ! ” --as used in many common programming languages--to indicate logical negation would preserve the original meaning of the tilde in case it becomes necessary in a future notation.
○ ***Recommendation 2*** – “*And*.” Similarly, the field chose “ & ” to represent the conjunction relationship between constraints. We recommended preserving this symbol use since it is human- and computer-readable and does not overlap with any notation in the original Linear Code.
○ ***Recommendation 3*** – “*Number*.” “ # ” was defined to combine glycans with glycosides other than amino acids and lipids (Table 3: GR1). Krambeck et al. uses it to represent the number of times a certain MS appears, a common use of “ # ”. In LiCoRR, we depreciate the use of “ # ”, “ ; ”, and “ : ” to specify the glycoconjugate class. The number sight “#” can be used to separate a glycan (on the left) from any conjugate (on the right). Colon and semi-colon can therefore be reserved for other future uses. To specify a glycopeptide, users may also inscribe them directly in the peptide using the existing branching rules: “PEP(AG(LY)CAN)TIDE” would describe a biantennary glycan bound to the Threonine of a peptide. Because the number sign is used to indicate a glycoconjugate, we recommend using “n.” For example, “#A = 5” will then be written as “nA=5” (Table 10: i).
○ ***Recommendation 4*** – “*Splitting & ‘or’*.” In addition to having several constraints split by “or,” we can rewrite the rules several times with a single constraint for each rule, as done for the reaction rule b4GalT in^14^. Splitting disjunctions over multiple lines is similar to atomization, the first normal form of database normalization requiring the domain of each attribute to contain an indivisible element. In addition, the separate rules have the advantage that they can have different reaction rate parameters. This advantage can eliminate the need for separate adjustment rules for the various cases. Depending on the circumstances, splitting disjunctions across multiple lines may be necessary, though it is often more succinct to condense them, separated by an “or” within a single rule.
○ ***Recommendation 5*** – “*Branch point*.” Many studies using Linear Code to define glycan synthesis networks assigned “ | ” as a possible branch point^7,10,13^. Our recommendation, however, is to use “ + ” instead of “ | ” as the branch point because “ | ” is already assigned within the original Linear Code. Additionally, we think “ + ” is more morphologically close to a branch.
○ ***Recommendation 6*** – “*Omission*.” Though “ * ” has been widely used by the systems glycobiology field to represent the reaction site, the original Linear Code rules actually specify “ * ” to stand for the omission of an entire saccharide unit (Table 3: UR3). We wish to minimize this inconsistency with the original Linear Code paper (Banin et al. 2002) as much as possible. Therefore, for “ * ”, we recommend preserving the meaning of the omission of one entire SU. In theory, according to Banin’s definition, a saccharide unit can be specified as “ ??? ”. Using “ * ” to indicate a complete SU, would avoid using an unmanageable number of question marks to represent an ambiguous glycan. Question marks should still be used to indicate unknown elements of an SU (e.g. “Ab4Gb” without knowledge of “b4G” could be written as “A???b”), but there should never be four adjacent question marks. We propose a substitute for reaction site in Recommendation 7.
○ ***Recommendation 7*** – “*Reaction site*.” The reaction site is the location of the first change to the glycan expression. Because “ * “ is already defined within Linear Code to indicate “omission,” we choose “ @ ” to indicate the reaction site. The reaction site, in previous reaction rules as “ * “ and going forward as “ @ “, is the position of the first difference between product and substrate strings in the Linear Code form.
○ ***Recommendation 8*** - *“Modification.”* As specified in the original Linear Code, we recommend using “[]” to represent known modifications (Table 3: MR1). For example, “A[2P]” represents a galactose with its second position modified by a phosphate. However, this specific modification may not always be known. Therefore, in addition to “[]” as exact modifications, we recommended using the “ $ ” sign to represent a possible modification site. For example, “A$GN” represents a GlcNAc connected to a galactose that might be modified.
○ ***Recommendation 9 -*** “Branching index.” In LiCoRR we have depreciated the original linear code branching rules due to redundancy and default to a version of BR1: *Reguardless of whether the MS are equivalnt,* the MS linked to the higher index carbon will branch (appear first in the written representation when read right to left, reducing to non-reducing end). This rule can be extended to glycopeptides providing a means of representing glycans directly embedded in a glycopeptide. “PEP(Gal[3S]b3(GNb6)AN)TIDE” would describe a trisaccharide O-glycan bound to the Threonine of a suspiciously named glycoprotein.

**Table 10.**
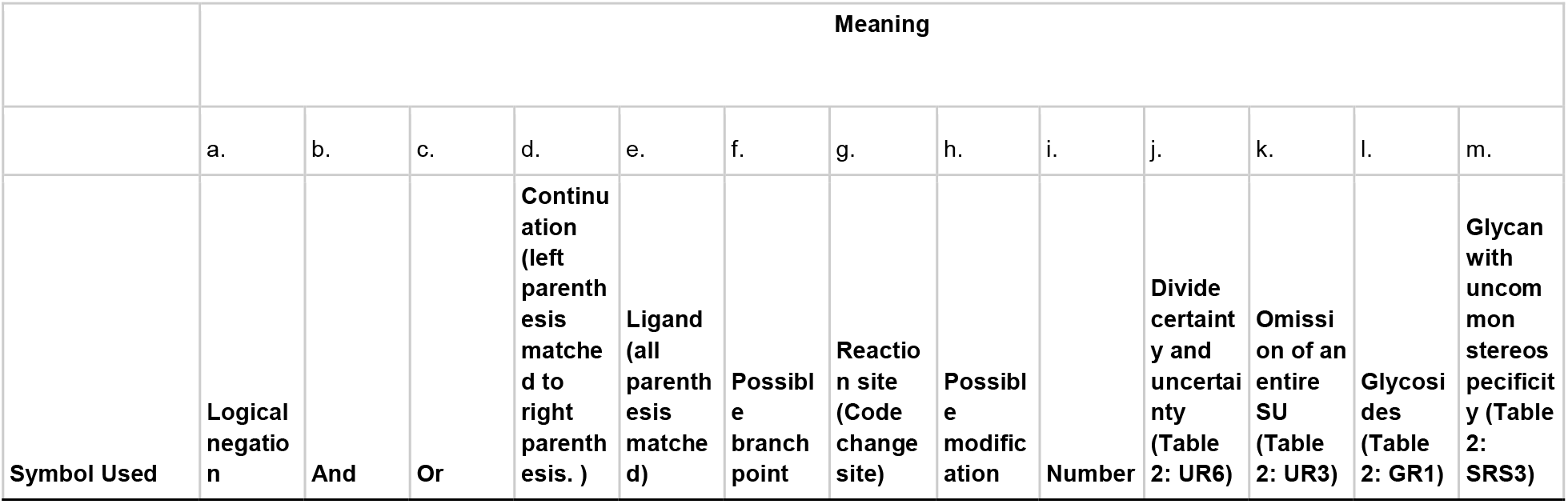
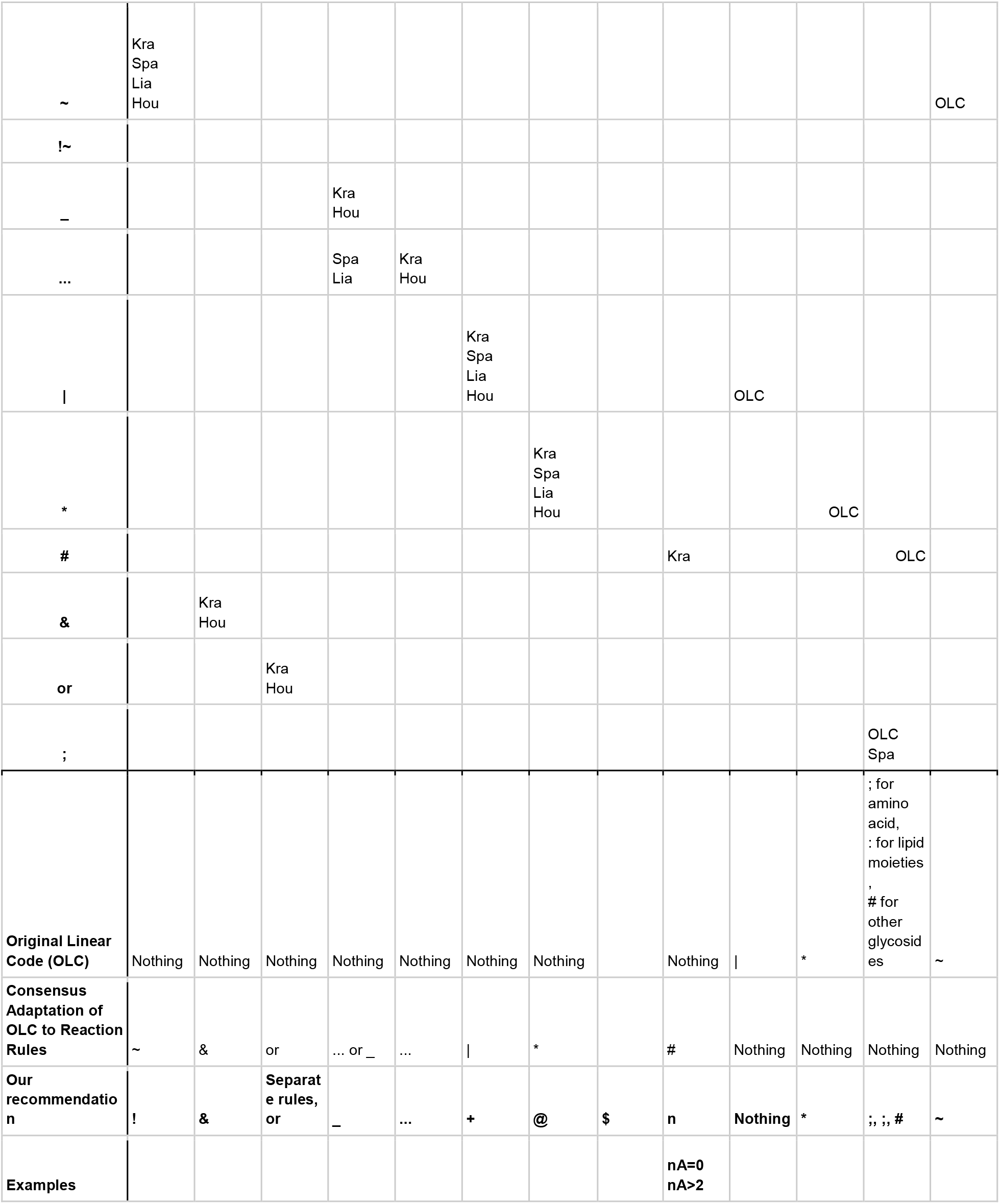
Symbols previously used by systems glycobiologists and our recommendations. Rows a - i are the functions implemented by published papers. Rows j - m are the functions prescribed in the original Linear Code rules.

Overall, the consensus in these representations centers around the foundational work of the original Linear Code paper^29^ and Krambeck et al. We have simply highlighted gaps in clarity that have resulted in colloquially small but computationally important divergences throughout the literature.

## Conclusion

The field of systems glycobiology is poised to tackle increasingly complex glycan synthesis problems owing to the advent of a number of enabling computational modeling technologies. Linear Code is used to represent reaction rules of glycan synthesis thereby bringing both human-readability and computer-parsability to the glycoinformatics space. The utility of Linear Code in glycoinformatics has been extended by the inclusion of new symbols, relations, and attributes that accommodate the challenge of specifying reaction rules. Yet various implementations conflict with each other and the original Linear Code. Here, we delineate the various adaptations made to accommodate reaction rule representation, the discordance between various implementations, and propose a consensus for future representations called LiCoRR.

The adoption of a common reaction rule representation would increase FAIR (Findable, Accessible, Interoperable, Reusable) standards^38^ compliance in glycoinformatics which will have far-reaching implications. With a predictable representation, reaction rules could be searched directly making them “findable.” Importantly, unifying the reaction rule nomenclature will make it possible to compare the performance and predictions of multiple models. Ideally, reaction rules sets will be sufficiently standardized so that they will be “interoperable” across multiple modeling software so that models can be “reused,” reproduced, validated and extended across labs. Increased readability and FAIRness through clarifying the nomenclature will help advance glycoinformatics technologies by making possible cross-platform and multi-omics integration and interpretation; interoperability may be enhanced through a community-endorsed vocabulary.

We further hope that the symbols described in this work, specifically the wildcards, will be used in other glycan representations. Definition of glycan classes can be useful for efficiently and unambiguously describing the key elements of large unruly glycans while only communicating the central information. Adoption of these symbols, now well defined symbols, by more popular representations, such as IUPAC, could increase both the flexibility and succinctness of those representations. We hope to encourage that adoption through our LiCoRRICE exemplars.

Increased FAIRness will facilitate the validation and distribution of developing glycoinformatics toolkits. Easy-to-use glycoinformatics toolkits, made possible by the fluency of interoperability across tools, are one mechanism by which glycobiology can be shared with the broader community of biology.

## Acknowledgments

This work was supported by generous funding from the Novo Nordisk Foundation provided to the Center for Biosustainability at the Technical University of Denmark (NNF10CC1016517) and from NIGMS (R35 GM119850). Additional funding was provided by the Triton Research & Experiential Learning Scholars (TRELS) program at the University of California, San Diego, the NIDDK Information Network (dkNET) supported by NIH & NIDDK (2U24DK097771-06), and NIH (NIH HL103411).

---

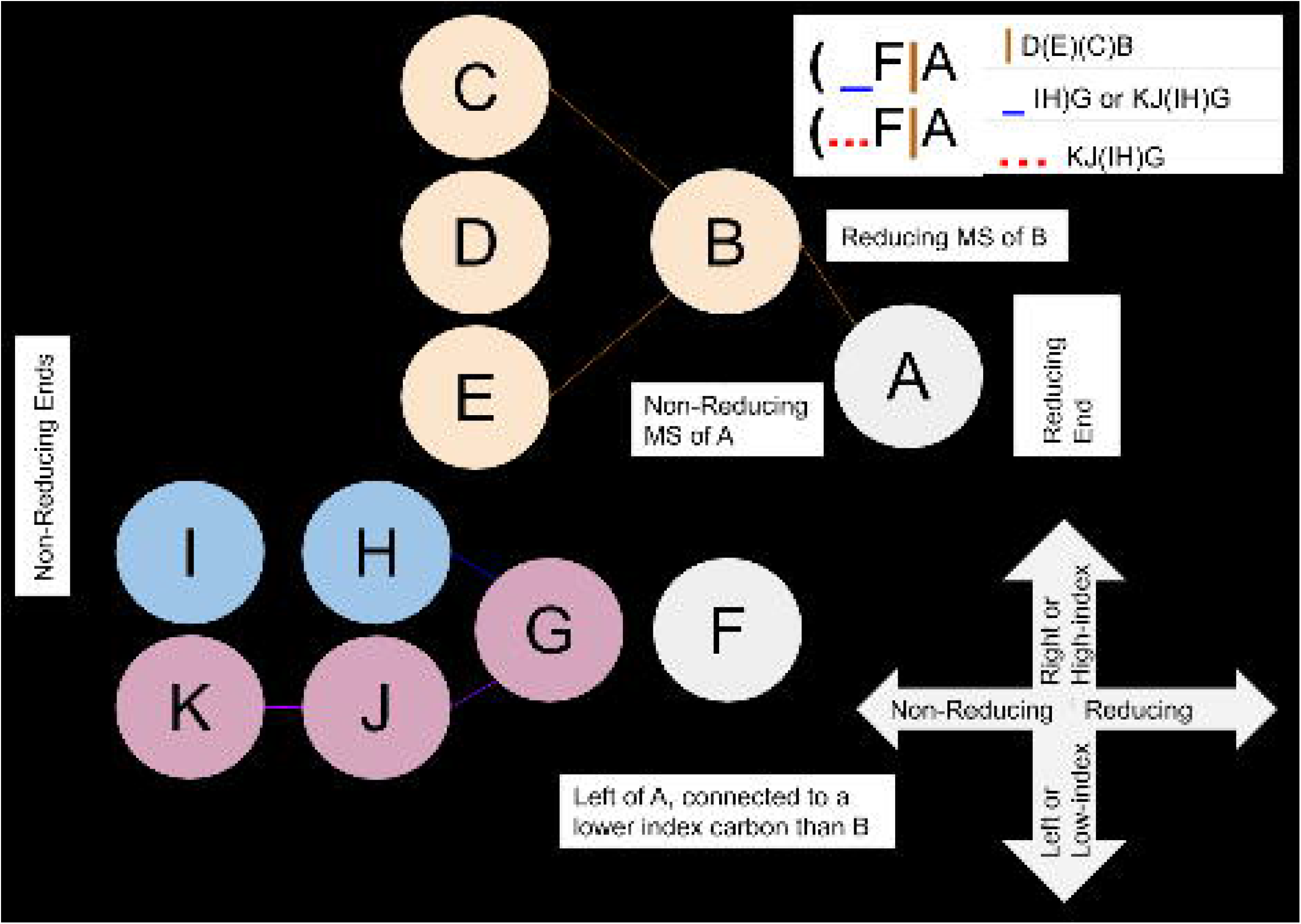

## Notes

### Competing Interest Statement

The authors have declared no competing interest.

## References

(1) Kontoravdi, C.; Jimenez del Val, I. Computational Tools for Predicting and Controlling the Glycosylation of Biopharmaceuticals. Curr. Opin. Chem. Eng. 2018, 22, 89–97.

(2) Spahn, P. N.; Lewis, N. E. Systems Glycobiology for Glycoengineering. Curr. Opin. Biotechnol. 2014, 30, 218–224.

(3) Puri, A.; Neelamegham, S. Understanding Glycomechanics Using Mathematical Modeling: A Review of Current Approaches to Simulate Cellular Glycosylation Reaction Networks. Ann. Biomed. Eng. 2012, 40 (4), 816–827.

(4) Krambeck, F. J.; Bennun, S. V.; Andersen, M. R.; Betenbaugh, M. J. Model-Based Analysis of N-Glycosylation in Chinese Hamster Ovary Cells. PLoS One 2017, 12 (5), e0175376.

(5) Sou, S. N.; Jedrzejewski, P. M.; Lee, K.; Sellick, C.; Polizzi, K. M.; Kontoravdi, C. Model-Based Investigation of Intracellular Processes Determining Antibody Fc-Glycosylation under Mild Hypothermia. Biotechnol. Bioeng. 2017, 114 (7), 1570–1582.

(6) Del Val, I. J.; Polizzi, K. M.; Kontoravdi, C. A Theoretical Estimate for Nucleotide Sugar Demand towards Chinese Hamster Ovary Cellular Glycosylation. Sci. Rep. 2016, 6, 28547.

(7) Hou, W.; Qiu, Y.; Hashimoto, N.; Ching, W.-K.; Aoki-Kinoshita, K. F. A Systematic Framework to Derive N-Glycan Biosynthesis Process and the Automated Construction of Glycosylation Networks. BMC Bioinformatics 2016, 17 Suppl 7, 240.

(8) Liu, G.; Neelamegham, S. Integration of Systems Glycobiology with Bioinformatics Toolboxes, Glycoinformatics Resources, and Glycoproteomics Data. Wiley Interdiscip. Rev. Syst. Biol. Med. 2015, 7 (4), 163–181.

(9) McDonald, A. G.; Hayes, J. M.; Bezak, T.; Głuchowska, S. A.; Cosgrave, E. F. J.; Struwe, W. B.; Stroop, C. J. M.; Kok, H.; van de Laar, T.; Rudd, P. M.; Tipton, K. F.; Davey, G. P. Galactosyltransferase 4 Is a Major Control Point for Glycan Branching in N-Linked Glycosylation. J. Cell Sci. 2014, 127 (Pt 23), 5014–5026.

(10) Krambeck, F. J.; Bennun, S. V.; Narang, S.; Choi, S.; Yarema, K. J.; Betenbaugh, M. J. A Mathematical Model to Derive N-Glycan Structures and Cellular Enzyme Activities from Mass Spectrometric Data. Glycobiology 2009, 19 (11), 1163–1175.

(11) Liu, G.; Marathe, D. D.; Matta, K. L.; Neelamegham, S. Systems-Level Modeling of Cellular Glycosylation Reaction Networks: O-Linked Glycan Formation on Natural Selectin Ligands. Bioinformatics 2008, 24 (23), 2740–2747.

(12) Liu, G.; Puri, A.; Neelamegham, S. Glycosylation Network Analysis Toolbox: A MATLAB-Based Environment for Systems Glycobiology. Bioinformatics 2013, 29 (3), 404–406.

(13) Spahn, P. N.; Hansen, A. H.; Hansen, H. G.; Arnsdorf, J.; Kildegaard, H. F.; Lewis, N. E. A Markov Chain Model for N-Linked Protein Glycosylation – towards a Low-Parameter Tool for Model-Driven Glycoengineering. Metabolic Engineering. 2016, pp 52–66. https://doi.org/10.1016/j.ymben.2015.10.007.

(14) Liang, C.; Chiang, A. W. T.; Hansen, A. H.; Arnsdorf, J.; Schoffelen, S.; Sorrentino, J. T.; Kellman, B. P.; Bao, B.; Voldborg, B. G.; Lewis, N. E. A Markov Model of Glycosylation Elucidates Isozyme Specificity and Glycosyltransferase Interactions for Glycoengineering. Current Research in Biotechnology 2020. https://doi.org/10.1016/j.crbiot.2020.01.001.

(15) Spahn, P. N.; Hansen, A. H.; Kol, S.; Voldborg, B. G.; Lewis, N. E. Predictive Glycoengineering of Biosimilars Using a Markov Chain Glycosylation Model. Biotechnol. J. 2017, 12 (2).

(16) McDonald, A. G.; Tipton, K. F.; Davey, G. P. A Knowledge-Based System for Display and Prediction of O-Glycosylation Network Behaviour in Response to Enzyme Knockouts. PLoS Comput. Biol. 2016, 12 (4), e1004844.

(17) Bao, B.; Kellman, B. P.; Chiang, A. W. T.; York, A. K.; Mohammad, M. A.; Haymond, M. W.; Bode, L.; Lewis, N. E. Correcting for Sparsity and Non-Independence in Glycomic Data through a Systems Biology Framework. bioRxiv. 2019. https://doi.org/10.1101/693507.

(18) Neelamegham, S.; Aoki-Kinoshita, K.; Bolton, E.; Frank, M.; Lisacek, F.; Lütteke, T.; O’Boyle, N.; Packer, N. H.; Stanley, P.; Toukach, P.; Varki, A.; Woods, R. J.; SNFG Discussion Group. Updates to the Symbol Nomenclature for Glycans Guidelines. Glycobiology 2019, 29 (9), 620–624.

(19) Mehta, A. Y.; Cummings, R. D. GlycoGlyph: A Glycan Visualizing, Drawing and Naming Application. Bioinformatics 2020. https://doi.org/10.1093/bioinformatics/btaa190.

(20) Cheng, K.; Zhou, Y.; Neelamegham, S. DrawGlycan-SNFG: A Robust Tool to Render Glycans and Glycopeptides with Fragmentation Information. Glycobiology 2017, 27 (3), 200–205.

(21) Herget, S.; Ranzinger, R.; Maass, K.; C-W v. GlycoCT—a Unifying Sequence Format for Carbohydrates. Carbohydrate Research. 2008, pp 2162–2171. https://doi.org/10.1016/j.carres.2008.03.011.

(22) Tanaka, K.; Aoki-Kinoshita, K. F.; Kotera, M.; Sawaki, H.; Tsuchiya, S.; Fujita, N.; Shikanai, T.; Kato, M.; Kawano, S.; Yamada, I.; Narimatsu, H. WURCS: The Web3 Unique Representation of Carbohydrate Structures. Journal of Chemical Information and Modeling. 2014, pp 1558–1566. https://doi.org/10.1021/ci400571e.

(23) Matsubara, M.; Aoki-Kinoshita, K. F.; Aoki, N. P.; Yamada, I.; Narimatsu, H. WURCS 2.0 Update To Encapsulate Ambiguous Carbohydrate Structures. J. Chem. Inf. Model. 2017, 57 (4), 632–637.

(24) Aoki-Kinoshita, K.; Agravat, S.; Aoki, N. P.; Arpinar, S.; Cummings, R. D.; Fujita, A.; Fujita, N.; Hart, G. M.; Haslam, S. M.; Kawasaki, T.; Matsubara, M.; Moreman, K. W.; Okuda, S.; Pierce, M.; Ranzinger, R.; Shikanai, T.; Shinmachi, D.; Solovieva, E.; Suzuki, Y.; Tsuchiya, S.; Yamada, I.; York, W. S.; Zaia, J.; Narimatsu, H. GlyTouCan 1.0--The International Glycan Structure Repository. Nucleic Acids Res. 2016, 44 (D1), D1237–D1242.

(25) Campbell, M. P.; Peterson, R.; Mariethoz, J.; Gasteiger, E.; Akune, Y.; Aoki-Kinoshita, K. F.; Lisacek, F.; Packer, N. H. UniCarbKB: Building a Knowledge Platform for Glycoproteomics. Nucleic Acids Res. 2014, 42 (Database issue), D215–D221.

(26) York, W. S.; Mazumder, R.; Ranzinger, R.; Edwards, N.; Kahsay, R.; Aoki-Kinoshita, K. F.; Campbell, M. P.; Cummings, R. D.; Feizi, T.; Martin, M.; Natale, D. A.; Packer, N. H.; Woods, R. J.; Agarwal, G.; Arpinar, S.; Bhat, S.; Blake, J.; Castro, L. J. G.; Fochtman, B.; Gildersleeve, J.; Goldman, R.; Holmes, X.; Jain, V.; Kulkarni, S.; Mahadik, R.; Mehta, A.; Mousavi, R.; Nakarakommula, S.; Navelkar, R.; Pattabiraman, N.; Pierce, M. J.; Ross, K.; Vasudev, P.; Vora, J.; Williamson, T.; Zhang, W. GlyGen: Computational and Informatics Resources for Glycoscience. Glycobiology 2019. https://doi.org/10.1093/glycob/cwz080.

(27) Alocci, D.; Mariethoz, J.; Gastaldello, A.; Gasteiger, E.; Karlsson, N. G.; Kolarich, D.; Packer, N. H.; Lisacek, F. GlyConnect: Glycoproteomics Goes Visual, Interactive, and Analytical. J. Proteome Res. 2019, 18 (2), 664–677.

(28) Panico, R.; Richer, J.-C. A Guide to IUPAC Nomenclature of Organic Compounds: Recommendations 1993 : (including Revisions, Published and Hitherto Unpublished, to the 1979 Edition of Nomenclature of Organic Chemistry); Blackwell Science Incorporated, 1995.

(29) Banin, E.; Neuberger, Y.; Altshuler, Y.; Halevi, A.; Inbar, O.; Nir, D.; Dukler, A. A Novel Linear Code® Nomenclature for Complex Carbohydrates. Trends Glycosci. Glycotechnol. 2002, 14 (77), 127–137.

(30) Bennun, S. V.; Yarema, K. J.; Betenbaugh, M. J.; Krambeck, F. J. Integration of the Transcriptome and Glycome for Identification of Glycan Cell Signatures. PLoS Comput. Biol. 2013, 9 (1), e1002813.

(31) Varki, A.; Sherman, W.; Kornfeld, S. Demonstration of the Enzymatic Mechanisms of α-N-Acetyl-D-Glucosamine-1-Phosphodiester N-Acetylglucosaminidase (formerly Called α-N-Acetylglucosaminylphosphodiesterase) and Lysosomal α-N-Acetylglucosaminidase. Arch. Biochem. Biophys. 1983, 222 (1), 145–149.

(32) Rohrer, J.; Kornfeld, R. Lysosomal Hydrolase Mannose 6-Phosphate Uncovering Enzyme Resides in the Trans-Golgi Network. Mol. Biol. Cell 2001, 12 (6), 1623–1631.

(33) Hare, N. J.; Lee, L. Y.; Loke, I.; Britton, W. J.; Saunders, B. M.; Thaysen-Andersen, M. Mycobacterium Tuberculosis Infection Manipulates the Glycosylation Machinery and the N-Glycoproteome of Human Macrophages and Their Microparticles. J. Proteome Res. 2017, 16 (1), 247–263.

(34) Röttger, S.; White, J.; Wandall, H. H.; Olivo, J. C.; Stark, A.; Bennett, E. P.; Whitehouse, C.; Berger, E. G.; Clausen, H.; Nilsson, T. Localization of Three Human Polypeptide GalNAc-Transferases in HeLa Cells Suggests Initiation of O-Linked Glycosylation throughout the Golgi Apparatus. J. Cell Sci. 1998, 111 (Pt 1), 45–60.

(35) Akune, Y.; Lin, C.-H.; Abrahams, J. L.; Zhang, J.; Packer, N. H.; Aoki-Kinoshita, K. F.; Campbell, M. P. Comprehensive Analysis of the N-Glycan Biosynthetic Pathway Using Bioinformatics to Generate UniCorn: A Theoretical N-Glycan Structure Database. Carbohydr. Res. 2016, 431, 56–63.

(36) McDonald, A. G.; Boyce, S.; Tipton, K. F. ExplorEnz: The Primary Source of the IUBMB Enzyme List. Nucleic Acids Res. 2009, 37 (Database issue), D593–D597.

(37) Agravat, S. B.; Song, X.; Rojsajjakul, T.; Cummings, R. D.; Smith, D. F. Computational Approaches to Define a Human Milk Metaglycome. Bioinformatics 2016, 32 (10), 1471–1478.

(38) Wilkinson, M. D.; Dumontier, M.; Aalbersberg, I. J. J.; Appleton, G.; Axton, M.; Baak, A.; Blomberg, N.; Boiten, J.-W.; da Silva Santos, L. B.; Bourne, P. E.; Bouwman, J.; Brookes, A. J.; Clark, T.; Crosas, M.; Dillo, I.; Dumon, O.; Edmunds, S.; Evelo, C. T.; Finkers, R.; Gonzalez-Beltran, A.; Gray, A. J. G.; Groth, P.; Goble, C.; Grethe, J. S.; Heringa, J.; ’t Hoen, P. A. C.; Hooft, R.; Kuhn, T.; Kok, R.; Kok, J.; Lusher, S. J.; Martone, M. E.; Mons, A.; Packer, A. L.; Persson, B.; Rocca-Serra, P.; Roos, M.; van Schaik, R.; Sansone, S.-A.; Schultes, E.; Sengstag, T.; Slater, T.; Strawn, G.; Swertz, M. A.; Thompson, M.; van der Lei, J.; van Mulligen, E.; Velterop, J.; Waagmeester, A.; Wittenburg, P.; Wolstencroft, K.; Zhao, J.; Mons, B. The FAIR Guiding Principles for Scientific Data Management and Stewardship. Sci Data 2016, 3, 160018.

