## Supplemental Text & Tables for "A consensus-based and readable extension of *Li*near *Co*de for *R*eaction *R*ules (LiCoRR)": 52820LiCoRR supporting information.docx

**Supporting Tables**	**3**

Table S1 - All position combination F(E(D)(C)B)A	3

Table S2 - All Reaction Rules Sets Examined in this manuscript	3

Table S3 - Wildcard Checker	3

Table S4 - Computational Illustration of Uncertainty Operator Matches	3

**Supporting Software**	**4**

gRegEx	4

Checking syntactic well-formedness	4

Converting linear code representation of a glycan to a readable textual tree representation	4

Checking whether a string matches an uncertainty operator in a glycan	5

Finding all nonempty subsequences that match an uncertainty operator in a glycan	5

A Context-free Grammar for LiCoRR representations	7

### Supporting Tables

#### Table S1 - All position combination F(E(D)(C)B)A

The glycan string F(E(D)(C)B)A is written in every valid form using all three wildcards. Each row shows the symbol “<>” in one position. The checkmark is shown in the next three columns if either a “...”, “_” or “+” can replace the “<>” to recapitulate the original string: F(E(D)(C)B)A. White MSs are certain MSs appearing in the linear code. Gray MSs are what the wild card symbol stands for in the F(E(D)(C)B)A glycan.

#### Table S2 - All Reaction Rules Sets Examined in this manuscript

All reaction rule sets discussed in the paper collected in LiCoRR format. These rules can be used for future analyses and multiple comparisons.

#### Table S3 - Wildcard Checker

An excel sheet with a symbol checker implemented in VBA. This program will check if substrings in the first column match either the continuation (“_”), ligand (“...”), or branch (“+”).

#### Table S4 - Computational Illustration of Uncertainty Operator Matches

The panels of this figure describe the (nonempty) subsequences of the richly structured glycan NNa3(ANb4)Ab4GNb2(NNa6Ab4GNb4)Ma3(NNa3(ANb4)Ab4GNb3Ab4GNb2(NNa3(ANb4)Ab4GNb6)Ma6)Ma4GNb4(Fa6)GN that match LiCoRR's ' + ' vs. '....' vs. '_' uncertainty operators. Table (A) enumerates every (nonempty) contiguous subsequence of the demonstration glycan, plus the left and right contexts of the subsequence, plus whether the center subsequence matches each of the uncertainty operators, or (finally) any of the uncertainty operators. Table (B) is a subset of Table (A) that includes only those rows where the center ('match') subsequence matches at least one uncertainty operator. Tables (C-E) enumerate the subsets of Table (A)'s first three columns where the center ('match') subsequence matches each of the three uncertainty operators in {'+','...', '_'}. Table (F) enumerates the subset of Table (A) where the center ('match') subsequence matches the continuation operator ('_') but *not* the ligand operator ('...'). All tables were calculated more or less directly from the gRegEx software library.

### Supporting Software: gRegEx

Code is available at <https://github.com/emeinhardt/gregex>.

gRegEx is an open source Python library and command-line tool for working with linear code representations of glycans and checking what structures will match what uncertainty operators. For example, command-line functionality currently includes (1) checking whether a linear code representation is syntactically well-formed--matches the rules for a linear code or LiCoRR expression, (2) converting a linear code representation to an equivalent textual representation (common in computational linguistics) that makes branching structure of longer glycans easier to see at a glance, (3) identifying all subsequences of a glycan (along with associated left- and right-contexts) that match a given uncertainty operator, and (4) checking whether something can be substituted for a particular uncertainty operator in a glycan containing that operator.

While the Python module supports composable and extensible manipulation of linear code representations, four key functions are available via the command-line and illustrated below. (More details are available via python -m gregex -h and in the repository documentation.)

#### Checking syntactic well-formedness

If passed a linear code expression representing a single glycan, a Boolean is returned indicating whether the expression can be parsed according to the default grammar (see further below).

```
$ python -m gregex ‘Ma6(Ma4)M’  #is ‘Ma6(Ma4)M’ well-formed?
True
```

#### Converting linear code representation of a glycan to a readable textual tree representation

While linear code will be more compact (on average) than more general tree notations when chains (‘unary branching’) is more typical than multi-child branching, the ‘bushier’ a glycan is and the longer a glycan is, the harder it will be for a human to see hierarchical structure at a glance and the more likely they are to make mistakes while reading or editing. ‘s-expressions’ (see below) are a textual notation for (potentially nested) lists taken from the Lisp family of programming languages and sometimes used in computational linguistics for representing syntactic trees. They remain machine readable while making tree structure more apparent, particularly when indented according to common conventions supported by common text editors.

$ python -m gregex 'NNa3(ANb4)Ab4GNb2(NNa6Ab4GNb4)Ma3(NNa3(ANb4)Ab4GNb3Ab4GNb2(NNa3(ANb4)Ab4GNb6)Ma6)Ma4GNb4(Fa6)GN' -e

(GN Fa6 (GNb4 (Ma4 (Ma6 (GNb6 (Ab4 ANb4 NNa3)) (GNb2 (Ab4 (GNb3 (Ab4 ANb4 NNa3))))) (Ma3 (GNb4 (Ab4 NNa6)) (GNb2 (Ab4 ANb4 NNa3))))))

When indented more or less according to common conventions, this becomes

```
(GN Fa6
    (GNb4 (Ma4 (Ma6 (GNb6 (Ab4 ANb4
                               NNa3))
                    (GNb2 (Ab4 (GNb3 (Ab4 ANb4
                                          NNa3)))))
               (Ma3 (GNb4 (Ab4 NNa6))
                    (GNb2 (Ab4 ANb4
                               NNa3))))))
```

Restoring the one-line representation is as simple as removing tab and newline characters from this multi-line string.

#### Checking whether a string matches an uncertainty operator in a glycan

Given a linear code representation with exactly one uncertainty operator in it, users can check whether a particular string can be substituted for the operator:

$ python -m gregex ‘Ma4_M’ -s ‘(Ma6)’ # does ‘(Ma6)’ match ‘_’ and yield a well-formed glycan?

True

$ python -m gregex ‘Ma4_M’ -s ‘(Ma6’

False

#### Finding all nonempty subsequences that match an uncertainty operator in a glycan

Given a linear code representation of a single glycan, the gRegEx command-line interface supports enumerating all subsequences that match a particular uncertainty operator:

$ python -m gregex 'Ma4(Ma6)M' -o '_'

(Ma6)

(Ma6)M

)

)M

M

Ma4

Ma4(Ma6)

Ma4(Ma6)M

Ma6

Ma6)

Ma6)M

Optionally including left and right context as well:

```
$ python -m gregex 'Ma4(Ma6)M' -o '_' -c | csvtk tab2csv -H -t | csvtk add-header -n Left,Match,Right | csvtk csv2md
Left    |Match    |Right
:-------|:--------|:-----
        |Ma4      |(Ma6)M
        |Ma4(Ma6) |M
        |Ma4(Ma6)M|
Ma4     |(Ma6)    |M
Ma4     |(Ma6)M   |
Ma4(    |Ma6      |)M
Ma4(    |Ma6)     |M
Ma4(    |Ma6)M    |
Ma4(Ma6 |)        |M
Ma4(Ma6 |)M       |
Ma4(Ma6)|M        |
```

A variant of this command can be used to identify all nonempty subsequences that both match a particular uncertainty operator and where a particular substring could be successfully substituted for that operator:

```
$ python -m gregex 'Ma4(Ma6)M' -o '_' -s '(Ma2)' -c | csvtk tab2csv -H -t | csvtk add-header -n Left,Operator_Match,Right,Valid_Substitution | csvtk csv2md
Left    |Operator_Match|Right |Valid_Substitution
:-------|:-------------|:-----|:-----------------
        |Ma4           |(Ma6)M|True
        |Ma4(Ma6)      |M     |True
        |Ma4(Ma6)M     |      |True
Ma4     |(Ma6)         |M     |True
Ma4     |(Ma6)M        |      |True
Ma4(    |Ma6           |)M    |True
Ma4(    |Ma6)          |M     |False
Ma4(    |Ma6)M         |      |False
Ma4(Ma6 |)             |M     |False
Ma4(Ma6 |)M            |      |False
Ma4(Ma6)|M             |      |True
```

#### A Context-free Grammar for LiCoRR representations

In addition to these practical functions, the software and documentation offer explicit (and in some cases, mathematically precise) specification of LiCoRR notation and operators. For example, the set of syntactically well-formed linear code representations of glycans accepted by the parsing functionality is specified by a context-free grammar. This is a declarative, unambiguous, and mathematical reference description of LiCoRR notation that is independent of any particular parser implementation (recall the difficulties described at the end of “Current Usage of Linear Code to represent Reaction Rules” in the main text). The current grammar currently supports almost all features of LiCoRR relevant to describing a complete glycan or sets of them (as opposed to fragments of a glycan or components of a reaction rule). Below is a simplified version of the grammar (meant for exposition) that describes LiCoRR expressions that pick out (at most) a single glycan:

expression ⟶ (subexpression branch+)? reducing_chain | λ

reducing_chain ⟶ complete_SU* reducing_SU

branch ⟶ "(" subexpression ")"

subexpression ⟶ (subexpression branch+)? nonreducing_chain

nonreducing_chain ⟶ complete_SU+

complete_SU ⟶ MS variant_information? modification? bond_anomericity bond_location

reducing_SU ⟶ MS variant_information? modification? bond_anomericity?

variant_information ⟶ "'" | "^" | "~"

modification ⟶ "[" bond_location? non_carbohydrate_moiety ("," bond_location non_carbohydrate_moiety)* "]"

non_carbohydrate_moiety ⟶ "Q" | "PE" | "IN" | "ME" | "N" | "T" | "P" | "PC" | "PYR" | "S" | "SH" | "EP"

bond_anomericity ⟶ "a" | "b"

bond_location ⟶ nonzero_digit

MS ⟶ "A" | "AN" | "B" | "E" | "F" | "G" | "GN" | "H" | "I" | "K" | "L" | "M" | "NG" | "NJ" | "NN" | "O" | "P" | "PH" | "R" | "S" | "U" | "W" | "X"

nonzero_digit ⟶ "1" | "2" | "3" | "4" | "5" | "6" | "7" | "8" | "9"

Here “⟶”, “|”, “(”, “)”, “?”, “+”, “*”, and “λ” have their usual formal-language theoretic meaning (see any introductory material on formal language and automata theory) as metalinguistic symbols (“is”, “or”, parentheses indicating grouping, “zero or one”, “one or more”, “zero or more”, and the unique empty string of length 0, respectively), all terminal symbols are quoted (except for the empty string λ), the choice of non-carbohydrate moiety symbols was drawn from Banin et al. (2002), and the choice of monosaccharide symbols was drawn from a mix of Banin et al. (2002) and Krambeck et al. (2009). Unlike the actual grammar underlying the currently implemented parser, the grammar above does not support any uncertainty operators at all - hence why the LiCoRR expressions it recognizes or generates only describe at most a single glycan. For simplicity of exposition, the grammar above also omits (unlike the actual grammar underlying the parser) repeated element notation. Neither the grammar above nor the currently implemented grammar support glycoconjugate notation or notation for multiple uncertain connection sites. Neither grammar enforces conventions about ordering of children or constraints about what is physically impossible - e.g. having two distinct monosaccharides bind at the same index number of a common ‘parent’ monosaccharide. Restrictions of the first kind are simply (syntactic) conventions and restrictions of the second kind are part of the *denotational semantics* of LiCoRR expressions as a formal language, not the syntax.

Major features for future releases include

1. parsing fragments of glycans in LiCoRR (e.g. recognizing ‘(Ma2)(Ma4)’ as a sequence of subtrees whose roots are siblings),
2. converting between such fragments and s-expressions,
3. adding support for the denotational semantics of linear code and “normal forms” that enforce conventions like child ordering based on bond index,
4. calculating the results of applying a LiCoRR reaction rule to a substrate or group of substrates,
5. calculating the set of products of a reaction network,
6. given two of a substrate class, a product class, or a constraint, calculate possible values for the third,
7. given a set of substrates and a set of products, reduce those to a (set of) substrate class(es) and a (set of) product class(es),
8. given a set of substrates, a set of products, and a reaction network, infer a probability distribution over paths that would transform the substrates into the products.
