## Supplementary figures and images for "A consensus-based and readable extension of *Li*near *Co*de for *R*eaction *R*ules (LiCoRR)"

### Context.pdf

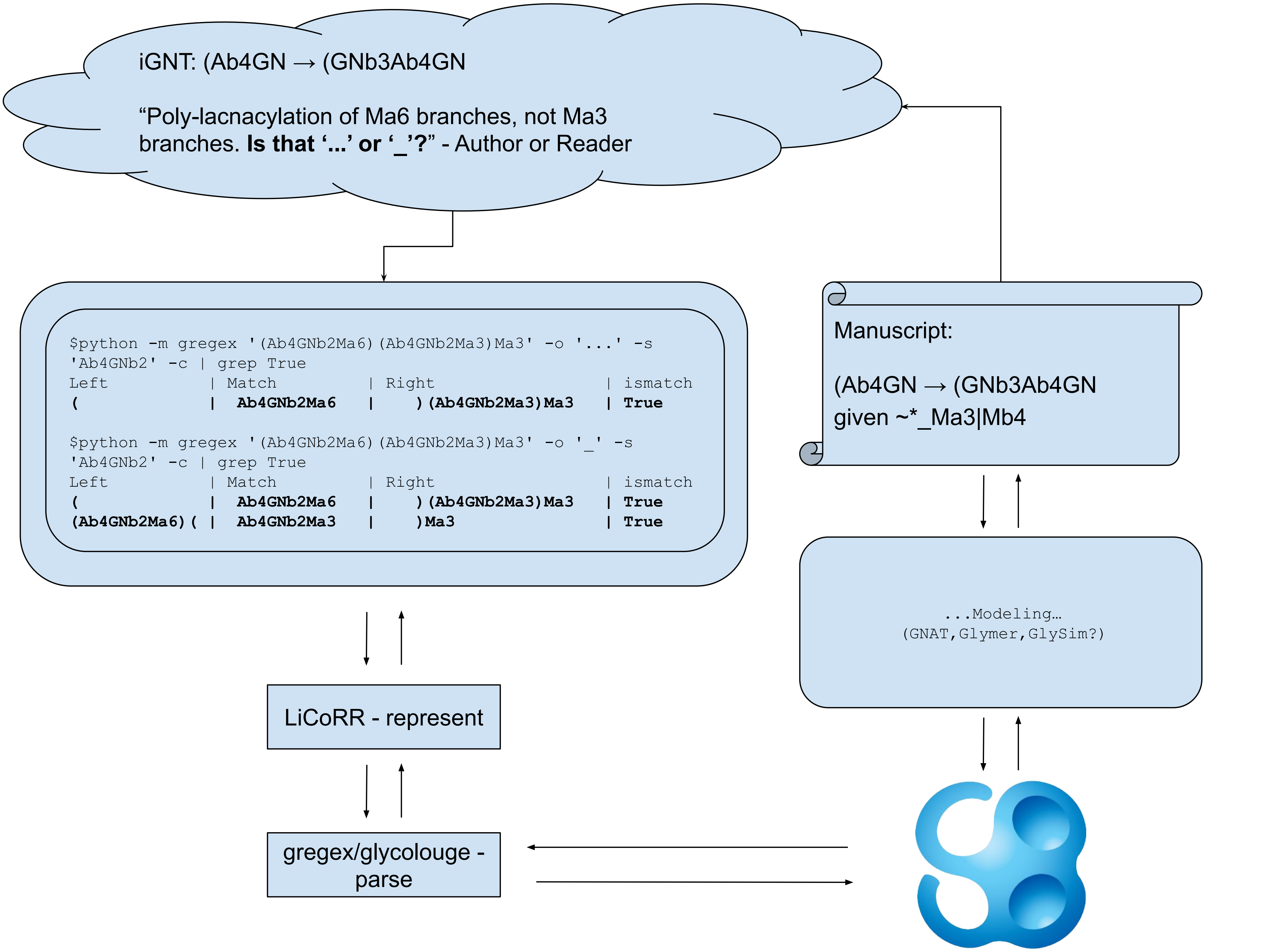

### fig1.pdf

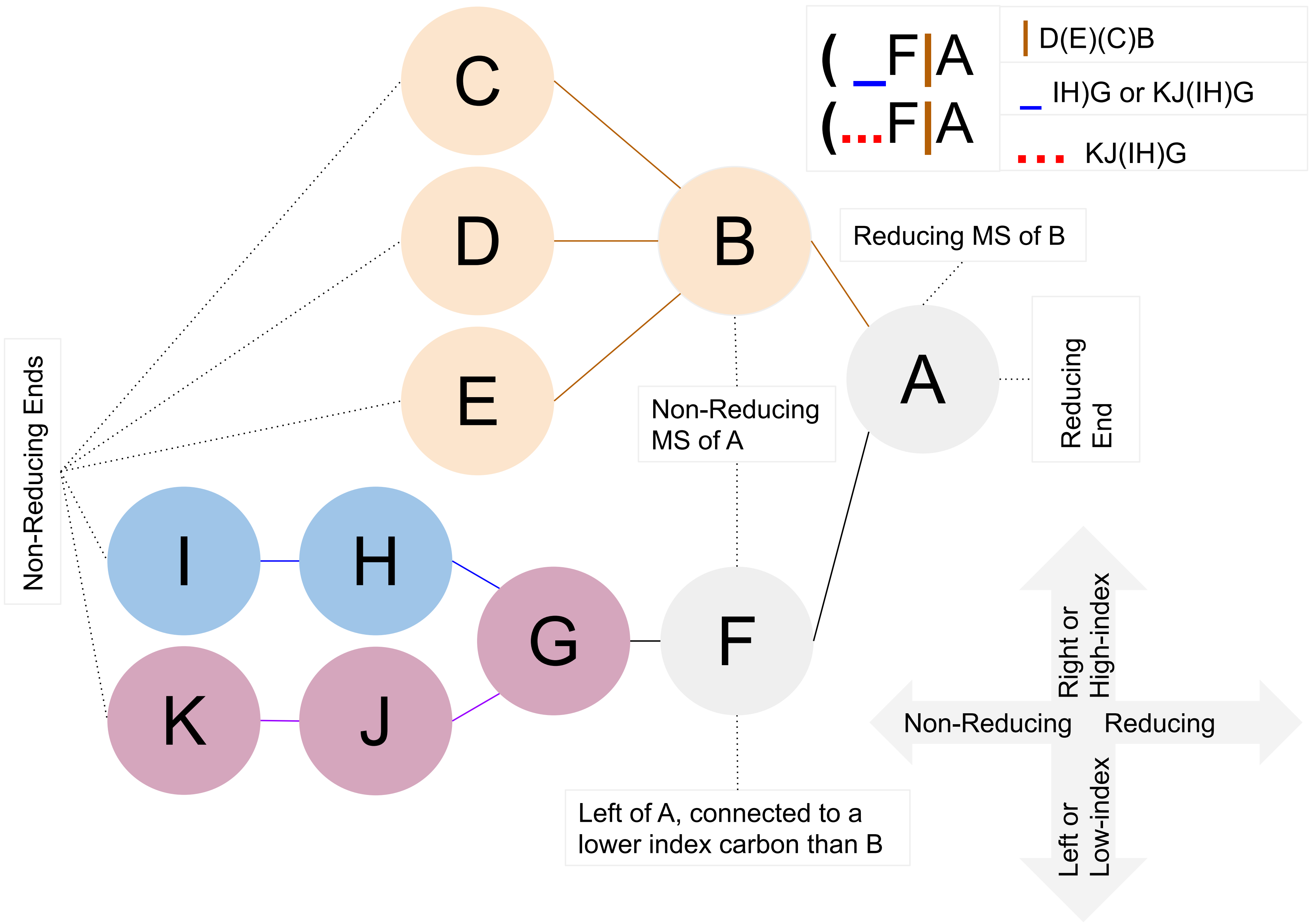
